# Integrated Metabolomic and Transcriptomic Profiling Reveals Novel Activation-Induced Metabolic Networks in Human T cells

**DOI:** 10.1101/635789

**Authors:** S. Hiemer, S. Jatav, J. Jussif, J. Alley, S. Lathwal, M. Piotrowski, J. Janiszewski, R. Kibbey, T. Alves, D. Dumlao, A. Jha, H. Bandukwala

## Abstract

The targeting of metabolic pathways is emerging as an exciting new approach for modulating immune cell function and polarization states. In this study, carbon tracing and systems biology approaches integrating metabolomic and transcriptomic profiling data were used to identify adaptations in human T cell metabolism important for fueling pro-inflammatory T cell function. Results of this study demonstrate that T cell receptor (TCR) stimulation leads to a significant increase in glucose and amino acid metabolism that trigger downstream biosynthetic processes. Specifically, increased expression of several enzymes such as CTPS1, IL4I1, and ASL results in the reprogramming of amino acid metabolism. Additionally, the strength of TCR signaling resulted in different metabolic enzymes utilized by T cells to facilitate similar biochemical endpoints. Furthermore, this study shows that cyclosporine represses the pathways involved in amino acid and glucose metabolism, providing novel insights on the immunosuppressive mechanisms of this drug. To explore the implications of the findings of this study in clinical settings, conventional immunosuppressants were tested in combination with drugs that target metabolic pathways. Results showed that such combinations increased efficacy of conventional immunosuppressants. Overall, the results of this study provide a comprehensive resource for identifying metabolic targets for novel combinatorial regimens in the treatment of intractable immune diseases.

## Introduction

Immune cells undergo profound metabolic changes during immune-cell activation to help them prepare and adapt for the increased anabolism required to sustain explosive proliferation and production of large amounts of biomolecules such as cytokines and immunoglobulins^1^. Wang et al. identified a key role for the transcription factor Myc in such metabolic reprogramming of T cells by showing that specific ablation of Myc suppressed pro-inflammatory functions in T cells^2^. Taken together, these studies support the hypothesis that targeting metabolic pathways in T cells could provide a novel therapeutic approach for the treatment of autoimmune and inflammatory diseases, thus resulting in a renaissance in the field of immunometabolism.

In addition to the role of immune cell metabolism in immune suppression and activation, recent reports have also revealed that specific metabolic pathways can also be used to regulate the fate and polarization state of immune cells. Rathmell and colleagues demonstrated that the induction of aerobic glycolysis upon TCR stimulation was critical for the pro-inflammatory functions of T cells. Blocking glucose uptake through ablation of the glucose transporter, Glut1, or altering the flux of pyruvate into the TCA cycle through modulation of pyruvate dehydrogenase activity were found to impact the inflammatory functions of T cells^3, 4^. In agreement with these observations Chi and colleagues identified HIF1α as a key transcription factor required for the increased aerobic glycolysis and pro-inflammatory functions of T cells^5^. Interestingly, blocking glycolysis not only suppressed differentiation of inflammatory Th17 cells but also promoted the differentiation of naïve T cells into Foxp3 expressing regulatory T cells. Got1 is another key enzyme that could regulate the polarization of both innate and adaptive immune cells. Got1 activity was selectively required for M2 macrophage polarization and for the differentiation of pathogenic Th17 cells. Additionally, inhibiting Got1 promotes the generation of Foxp3 regulatory T cells at the expense of Th17 differentiation^6^. These data provide further evidence that targeting key metabolic pathways can suppress the induction of inflammatory functions of immune cells as well as alter the polarization state of both innate and adaptive immune cells.

Pioneering works by several groups have highlighted the potential of targeting metabolic pathways not only for suppression of immune responses but also for augmenting immune cell function. Koichi and colleagues demonstrated that transient suppression of mTOR activation during priming of naïve T cells promotes the generation of memory T cells^7^. Pearce et al. similarly reported that administration of the anti-diabetic drug metformin promoted memory T cell differentiation through AMPK activation and indirect inhibition of mTOR function^8^. Corroborating these data, Rao and colleagues showed that mTOR inhibition promoted persistent Eomesodermin expression which in turn increased memory cell differentiation and recall responses^9^. Sukumar and colleagues translated these findings in pre-clinical oncology models and reported that inhibition of glycolysis during early T cell priming could enhance the memory and antitumor function of CD8+ T cells^10^. These data showed that modulation of immune cell metabolism could alter cellular differentiation programs. Additionally, these data highlighted that therapeutic applications of immunometabolism modulation are not limited to inflammatory diseases but can also be extended to anti-viral and immuno-oncology applications.

While there is increased enthusiasm to develop novel therapies based on advances in the understanding of immune cell metabolism, a majority of these studies have investigated metabolic programs in murine immune cells and not human cells. Further work is needed to successfully translate these results to a clinical setting and explore whether metabolic pathways that influence the function of murine immune cells are evolutionarily conserved in human immune cells. Hess and colleagues have demonstrated that some key observations obtained through the study of murine T cell metabolism are well conserved in human T cells^11–14^. Pearce and colleagues demonstrated a key ‘moonlighting’ role for the glycolytic enzyme Gapdh in murine T cells to suppress translation of IFN-γ by binding to the 3’-UTR of its mRNA. Increased aerobic glycolysis induced upon T cell activation reduces Gapdh binding to the IFN-γ UTR thus promoting its translation^15^. Hess and colleagues demonstrated that this function of Gapdh was evolutionarily conserved in human T cells where increased aerobic glycolysis was required for IFN-γ production by human effector T cells^11^. Furthermore, Geltink et al. demonstrated that CD28 costimulation promoted mitochondrial remodeling and increased spare respiratory capacity of murine CD8+ T cells. These processes were required for increased effector function in vitro and increased antitumor activity in vivo^16^. Bantug and colleagues demonstrated that similar mechanisms were utilized by human memory T cells which have expanded numbers of mitochondria in close approximation with the endoplasmic reticulum forming “Immunometabolic Hubs” that promoted their recall response^14^.

While these studies demonstrate that some key metabolic pathways are utilized similarly by both murine and human T cells, a few recent papers have highlighted some important differences in the human and murine immune system. These differences are in part due to the evolutionary divergence of these species 65 million years ago^17^ but also in part due to the highly regulated and specific pathogen-free conditions most laboratory mice are housed in. Masopoust and colleagues demonstrated that housing laboratory mice under environmental conditions more similar to what humans are exposed to led to significant changes in their immune systems^18^. These data suggest that there may be important differences by which human and murine immune cells may function due to differences in both environmental exposure as well as species-specific genetic architecture. A striking example of this difference was recently highlighted by Akondy et al. who demonstrated that quiescent human memory cells generated in response to vaccination had a doubling time which was over 450 days^19^. In sharp contrast, orthologous memory cells in mice generated in response to viral encounter had an intermitotic interval of only 50 days^7, 20, 21^. The 9-fold difference in the rates of turnover of human and murine memory T cells are likely to require different metabolic adaptations. To further explore differences and similarities between murine and human metabolism, we sought to profile the metabolic pathways that are triggered in human effector T cells upon TCR stimulation that help fuel their ability to proliferate and produce inflammatory cytokines.

In this study, we investigated the fate of two fundamental nutrients essential for T cell function, glucose and glutamine, to elucidate metabolic changes that are induced in primary human CD4+ T cells upon TCR stimulation. Using phenotypic assays, carbon tracing experiments, and a system-wide approach combining metabolomic and transcriptomic profiling, we show that human effector T cells primarily use glucose to fuel TCA anaplerosis but not mitochondrial respiration. The results of this study reveal a key adaptation of human effector T cells which allows them to undergo rapid proliferation while secreting energy expensive biomolecules like cytokines and chemokines. Additionally, we found that TCR stimulation of human effector T cells upregulates numerous metabolic enzymes which synergistically utilize amino acids such as glutamine and arginine to fuel TCA. Furthermore, analysis of integrated metabolomic and transcriptional profiling studies show that clinically relevant immunosuppressive drugs such as rapamycin and cyclosporine suppress the upregulation of amino acid metabolism pathways. These results provide new insights into the mechanisms by which these clinical agents limit inflammatory processes. Additionally, the results in this study reveal the suppressive effects of these clinical agents on additional metabolic pathways, which may in turn result in toxic adverse events of these agents. Finally, by combining CRISPR-based gene ablation and pharmacological inhibition studies, we dissect the contribution of TCR-induced signaling pathways to identify novel metabolic enzymes that could be targeted for more precise immune regulation in the absence of dose-limiting adverse events. The data and results of this study provides an important stepping-stone for the translation of exciting advances in immunometabolism into clinical therapies.

## Materials and Methods

### Primary human CD4+ T cell isolation, culture, and activation

Blood was collected from healthy human patients in accordance with IRB standards. CD4+ T cells were purified using RosetteSep isolation kit (StemCell) as per manufacturer’s protocol. Cells were cultured in RPMI supplemented with 10% heat inactivated FBS, HEPES, penicillin/streptomycin, non-essential amino acids, glutamax, and sodium pyruvate. Cells were initially activated for 72 hours on plate-bound α-CD3 (200 ng/mL; OKT3; BioXCell) and α-CD28 (200 ng/mL; 9.3; BioXCell) on goat-α-mouse (10 µg/mL; Jackson Immuno Research Labs) pre-coated high-binding plates (Corning Costar). After 72 hours, cells were removed from plate activation and were expanded until Day 8-12, maintained at ∼1X10^6^ cells/mL by supplementing with fresh media as required. Cells were then washed with PBS and either rested or re-activated at 0.5-1X10^6^ cells/mL in fresh media with plate-bound α-CD3/α-CD28 (100 ng/mL each) or with soluble ImmunoCult CD3/CD28 (1:100 dilution; StemCell). Cells were treated with Rapamycin (100 nM) and/or Cyclosporin A (2 µM), or equivalent volume of DMSO during re-activation.

### Lentiviral transduction and shRNA knockdown

Lenti-X 293T (Clontech) cells were plated on poly-D-lysine (0.1 mg/mL; Sigma) coated plates at a density of 1.67X10^5^ cells/mL. After 72 hours, cells were transfected with psPAX2 (1.5 µg), pCMV-VSV-G (0.5 µg), and pZIP-mCMV-ZsGreen-shRNA NT Control or mTOR plasmids (2 µg; Transomic) using the TransIT-LT1 transfection reagent (Mirus) as per manufacturer’s protocol. Virus was collected after an additional 72 hours and CD4+ T cells were spin infected after 48 hours of activation using undiluted virus (∼1X10^8^ IFU yielding a MOI of ∼70) supplemented with polybrene (10 µg/mL; AmericanBio Inc). After an additional 24 hours of activation, the transduced CD4+ T cells were removed from activation and expanded in the presence of puromycin (1 µg/mL) until Day 8-12, maintained at ∼1X10^6^ cells/mL by supplementing with fresh media as required. Cells were then washed and activated as described above.

### CRISPR/Cas-9 knockout

Cells were activated on plate-bound α-CD3/α-CD28 as described above. Cells were removed from plate activation after 72 hours and were rested overnight at ∼1X10^6^ cells/mL. The following day the conditioned media was saved before washing the cells and resuspending in buffer T (Neon Transfection System; ThermoFisher) at 25X10^6^ cells/mL. Guides targeting human mTOR and ZAP70 were designed using the Broad Institute’s online sgRNA designer (https://portals.broadinstitute.org/gpp/public/analysis-tools/sgrna-design)22 (see Sup Table 2). 5 guides per gene were synthesized by IDT and duplexed in a 1:1 ratio with Alt-R-CRISPR-Cas9-tracrRNA (IDT) as per manufacturer’s protocol. Duplexed guides were incubated with Alt-R S.p. Cas9 Nuclease (IDT) before mixing with 2.5X10^6^ cells and electroporating using the Neon Transfection System (ThermoFisher; 100uL tip at 1550V, 10 milliseconds, 3 pulses). Cells were allowed to recover for 1 hour in fresh media before adding back conditioned media in a 1:1 ratio conditioned to fresh media. Cells were expanded in the presence of IL-2 (2 ng/mL; R&D) until Day 8-12, maintained at ∼1X10^6^ cells/mL by supplementing with fresh media as required. Cells were then washed and activated as described above.

### Antibodies, Dyes, and Flow cytometry analysis

Cells were washed and stained with Live/Dead Fixable Near-IR Dead Cell stain (1:1000; ThermoFisher) for 10 min at room temperature. Cells were washed, fixed in 4% paraformaldehyde for 10 min at room temperature, and washed again for downstream staining or FACS analysis. Cells were permeabilized in 90% methanol for 1 hour at −20°C, washed, and stained overnight with α-human-phospho-S6(PS235/PS236)-AF647 (1:50; BD Biosciences) or α-human-ZAP70-PE (1:30; Biolegend) and equivalent isotype control. To assess mitochondrial mass and polarity, viable cells were incubated in MitoTracker Orange CM-H_2_TMRos (250 nM; ThermoFisher) for 30 min at 37°C. To assess ROS, viable cells were incubated in CellRox Orange (750 nM; ThermoFisher) for 1 hour at 37°C. To assess apoptosis, viable cells were incubated in Annexin V Pacific Blue (1:100; ThermoFisher) for 15 min at room temperature. After incubations, cells were washed, stained with viability stain, and fixed as described above. To assess proliferation, viable cells were incubated in CellTrace Violet (2 µM; ThermoFisher) for 20 min at 37°C. Cells were washed with FBS and fresh media and activated for 72-96 hours as described above. Cells were then harvested, stained with viability stain, and fixed as described above. Flow cytometry data were collected on a BD LSRFortessa (BD Biosciences) and analyzed using FlowJo software (TreeStar). 10000 events were collected, gated on the viable>singlet>ZsGreen positive population; or the viable>singlet population.

### Cytokine analysis

Supernatants were collected at the indicated time point and assessed using the V-PLEX Plus Proinflammatory Panel 1 (human) Kit (Meso Scale Diagnostics) according to manufacturer’s protocol.

### Metabolic Assays

ECAR and OCR were measured using a Seahorse XFe96 Analyzer (Agilent). Cells were counted and plated on the day of the experiment in assay media (Seahorse XF base medium supplemented with glutamine (2 mM), with or without sodium pyruvate (1 mM) or glucose (10 mM), adjusted to pH 7.4). Cells were plated at 0.4X10^6^ cells/well on Cell-Tak (22.4 µg/mL; Corning) coated plates. Final concentrations of the following compounds were used: Glucose (10 mM), Oligomycin (1 µM), 2-Deoxyglucose (50 mM), FCCP (500 nM), Rotenone+Antimycin A (1 µM), Etomoxir (40 µM), BPTES (3 µM), UK5099 (2 µM). Data was analyzed using Wave (Agilent) and GraphPad Prism software. Supernatants were removed immediately following the assay and lactate production was measured using the L-Lactate Assay Kit (Colorimetric; Abcam).

### RNA Sequencing

5X10^6^ cells were collected at the indicated time points, lysed with RNeasy (Qiagen) kit, and purified according to manufacturer’s protocol. RNA quantity was assessed using the QIAxpert and RNA quality was assessed using the Agilent 4200 TapeStation. RNA library preparation was performed using Illumina Stranded mRNA library preparation kit and sequencing was performed using NextSeq High Output Run with single end read 1×75bp.

### Untargeted metabolomics

300 μl of extraction buffer (acetonitrile:methanol:water 2:2:1) was added to frozen cell pellets (1X10^6^ cells) on ice and allowed to stand for 30 min. Samples were vortexed then centrifuged for 10 min at 15,000 RPM. The supernatants were transferred to new vial and dried under vacuum centrifugation. Samples were resuspended with 100 μl of resuspension buffer (acetonitrile:water 5:95). 10 μl of each sample was injected on our LC-HRMS system (Agilent 1290 UPLC in tandem with a Thermofisher High Field Q-Exactive) and separated using an ion-pairing chromatography.

#### Positive mode

Waters Acquity 2.1X100 mm BEH C18, 1.7μ column heated to 50°C. Mobile phase A – 5 mM perfluoropentanoic acid, mobile phase B – 50% Acetonitrile; flow rate: 0.4 mL/min; gradient – time 0 min = 2% B, time 0.5 = 2% B, time 10 min = 60% B, time 10.2 min = 90% B, time 12 min = 90% B, time 12.2 min = 50% B, time 14 min = 50% B, time 14.2 min = 2% B, time 18 min = 2% B. MS parameters: 120,000 resolution, AGC Target 1.0X10^6^, Maximum IT 100 ms, Sheath Gas Flow 30, Aux Gas Flow 5, Sweep Gas Flow 5, Spray Voltage (Kv) 4.3, Capillary Temp (°C) 275, S Lens RF 70, Aux Heat Temp (°C) 250.

#### Negative mode

Waters Acquity 2.1X100mm HSS T3, 1.7μ column heated to 35°C. Mobile phase A – 5% Methanol + 10 mM tributylamine & 15 mM acetic acid, mobile phase B – Isopropyl alcohol; flow rate: 0.4 mL/min; gradient – time 0 min = 0% B, time 4 = 0% B, time 7 min = 2% B, time 12.5 min = 10% B (flow rate 0.35 mL/min), time 13 min = 10% B (flow rate 0.3 mL/min), time 17 min = 25% B (flow rate 0.2 mL/min), time 20 min = 25% B (flow rate 0.2 mL/min), time 23 min = 0% B (flow rate 0.3 mL/min), time 24 min = 0% B (flow rate 0.4 mL/min), time 29 min = 0% B (flow rate 0.4 mL/min). MS parameters: 120,000 resolution, AGC Target 1.0X10^6^, Maximum IT 100 ms, Sheath Gas Flow 30, Aux Gas Flow 5, Sweep Gas Flow 5, Spray Voltage (Kv) 4.3, Capillary Temp (°C) 275, S Lens RF 70, Aux Heat Temp (°C) 250.

### Glutamine/Glucose labeling

Human CD4+ T cells were isolated and expanded as described above. On day 12 post initial activation, cells were either left unstimulated or acutely restimulated with ImmunoCult as described above. Cells were subsequently harvested and incubated in fresh cold media (RPMI with glucose or glutamine (MP Biomedicals) supplemented with sodium bicarbonate (3.7 g/L), HEPES (10 mM), essentially FFA free BSA (0.2%; Sigma), sodium pyruvate (0.05 mM), and L-lactate (0.45 mM; Sigma) for 1 hour. Subsequently cells were washed and pulsed with media containing labeled Glucose[U-^13^C_6_] (5.5 mM; CLM-1396; Cambridge Isotopes) or labeled Glutamine[1,2-^13^C_2_] (4 mM; CLM-2001; Cambridge Isotopes) for 4 hours. Cells were then washed with ice-cold PBS and immediately quenched with ice-cold 20% methanol in 0.1% formic acid supplemented with D4-taurine (10 µM; Cambridge Isotopes), Sodium Fluoride (3 mM; Sigma), phenylalanine (1 mM; Sigma), and EDTA (100 µM).

### MIMOSA Fluxomics

#### Extraction and Mass Spectrometry

Frozen quenched samples were lyophilized to dryness overnight in V-bottom plates (Bioexpress; Cat No. T-3025-8B). Samples were resuspended in water and 2 μL of each sample was analyzed on our LC-DMS/MS system (Agilent 1290 UPLC in tandem with Sciex 6500+ equipped with a DMS frontend). The method was previously described by Alves et al^23^. Briefly, glycolytic metabolites were analyzed using Imtakt amino acid 50 × 2.1 mm, 3μ column with the following mobile phase scheme: mobile phase A – 100 mM ammonium acetate, pH 6.9 and mobile phase B – acetonitrile. At a constant flow rate of 0.600 mL/min, metabolites were measured with the following gradient: time = 0 min (90% B), time = 0.75 min (90% B), time = 2 min (70% B), time = 3.6 min (70% B), time = 4 min (90% B), time = 5 min (90% B). The instrument parameters used are as follows: Curtain gas 30, CAD 9, IonSpray Voltage −3000, Temp 450, Gas 1 60, Gas 2 60, DMS Temp low, DMS Modifier 2-propanol, DMS Modifier Composition low, SV 2200, DR off, DMS Modifier flow (μL/min) 250, Dwell 10 msec, Pause 3 msec, Column Temp RT. TCA metabolites were analyzed using Hypercarb 100 × 2.1 mm column with the following mobile phase scheme: mobile phase A – 15 mM ammonium formate, 20 μL 0.5 EDTA pH 6.9 and mobile phase B – acetonitrile. At a constant flow rate of 0.400 mL/min, metabolites were measured with the following gradient: time = 0 min (0% B), time = 1 min (0% B), time = 1.75 min (1.5% B), time = 3.2 min (1.5% B), time = 3.5 min (0% B), time = 7 min (0% B). The instrument parameters used are as follows: Curtain gas 30, CAD 9, IonSpray Voltage −3000, Temp 450, Gas1 55, Gas 2 55, DMS Temp medium, DMS Modifier 2-propanol, DMS Modifier Composition low, SV 2600, DR open, DMS Modifier flow (μl/min) 325, Dwell 10 msec, Pause 3 msec, Column Temp RT. Optimized MRM and DMS parameters for each metabolites are detailed in Sup Table 3 (MRM).

### Data analyses

All mass spectrometry data was processed using Elucidata’s (www.elucidata.io) software packages: El-Maven and Polly.

#### Steady state precursor-product isotopic ratio (phi) analysis

The processing for MIMOSA fluxomics data was done through El-Maven. Briefly, peaks were picked for each of the MRM transitions (Sup Table 3) and peak areas were exported as a .csv file. PollyPhi^TM^ LC-MS/MS Relative app was then used to calculate ratios of various precursor and product fragments at isotopic steady state as described by Alves et al^23^. A phi value of 1 typically implies that there is no dilution of the isotopic label between the precursor and the product, whereas a ratio of less than 1 implies dilution due to either unlabeled carbons or labeled forms other than from the precursor. A description of the phi values used in this work along with the equations used to calculate them are provided in Sup Table 4.

#### Metabolomic data analysis

The processing for metabolomic datasets was done through El-Maven (https://doi.org/10.5281/zenodo.2537593). The compound database used for processing was derived from KEGG^44^ database, our final compound database consisted of all the compounds found in all of the KEGG’s reaction database. Automated peak picking feature of El-Maven was then used to get the peak intensity of all the metabolites. To get the differential expression between different cohorts metaboanalyst^45^ was used (http://www.metaboanalyst.ca). To annotate the metabolomic peaks, the mass which had least p-value in differential expression was selected. Enrichment of pathways in figure 3f was calculated using metaboanalyst.

**Figure 1:**
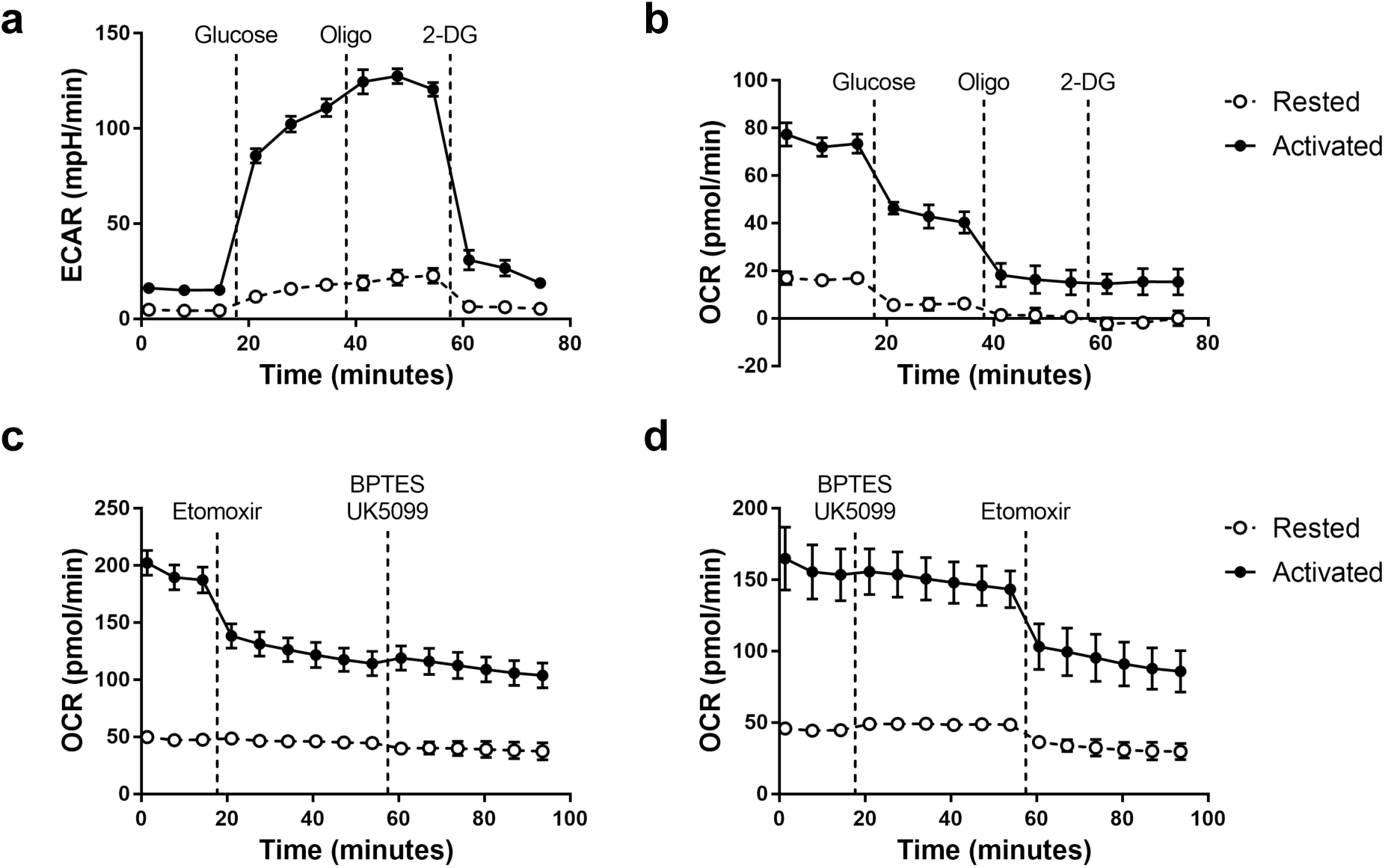
Utilization of glucose through MPC or glutamine through GLS1 are dispensable for mitochondrial ATP production in activated human CD4+ T cells. Primary human CD4+ T cells were rested or activated for 24 hours with CD3/CD28 ImmunoCult. a, ECAR and b, OCR measurements from one representative donor in response to glucose, oligomycin (Oligo), and 2-deoxyglucose (2-DG). OCR measurements from one representative donor in response to Etomoxir, BPTES, and UK5099 indicating fatty acid c, dependency and d, capacity.

**Figure 2:**
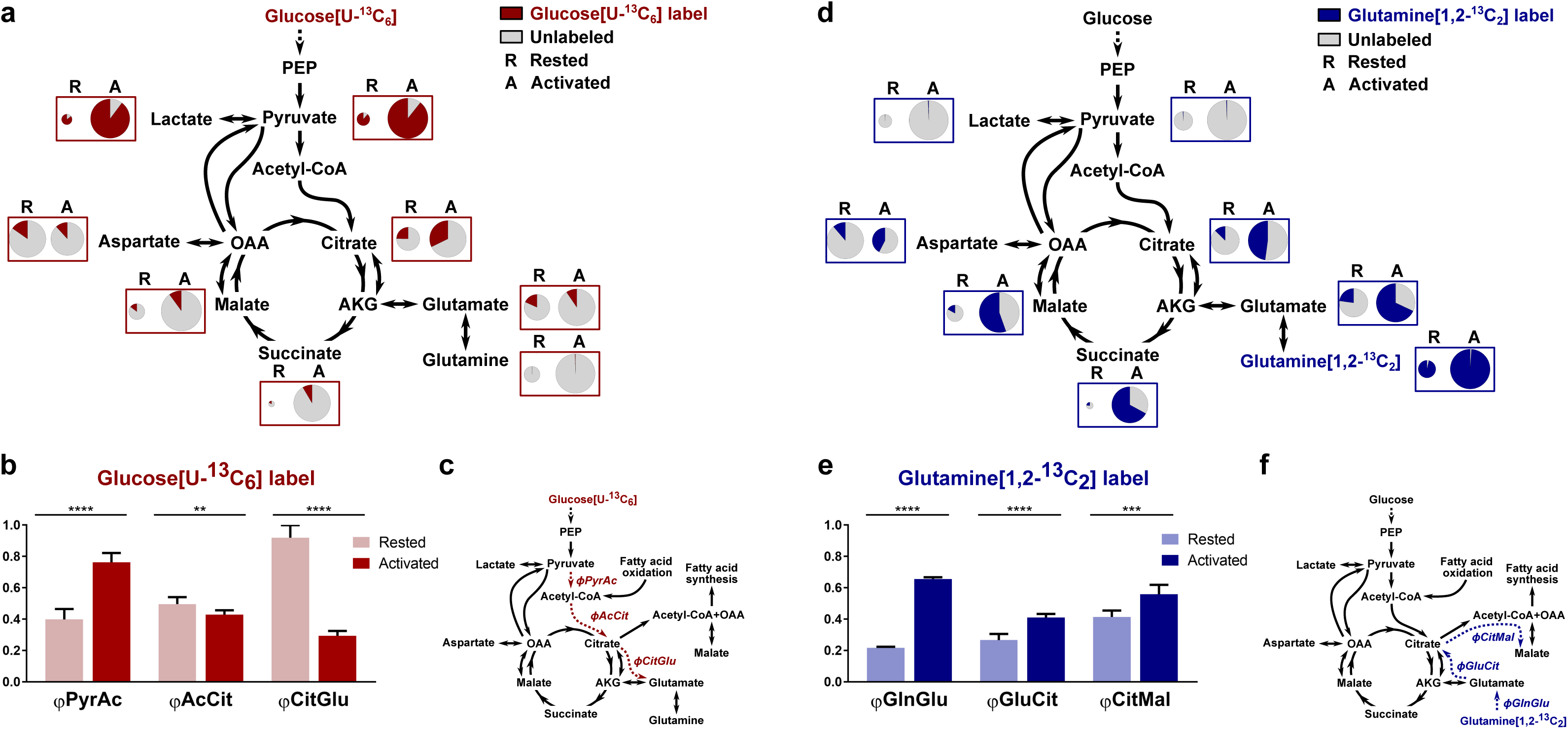
Glucose- and glutamine-derived carbon are used in anaplerotic processes in human CD4+ T cells. Primary human CD4+ T cells from one donor were rested or activated for 24 hours with CD3/CD28 ImmunoCult. a, Incorporation of [U-^13^C_6_]-D-glucose label (red) compared to unlabeled metabolites (grey) in Rested or Activated cells. Circle size corresponds to the pool size of each metabolite. b, Phi values of select metabolites. c, Schematic of glucose label flow. d, Incorporation of [1,2-^13^C_2_]-L-glutamine label (blue) compared to unlabeled metabolites (grey) in Rested or Activated cells. e, Phi values of select metabolites. f, Schematic of glutamine label flow. Data are presented as the mean ±SD, n=6. Unpaired two-tailed Student’s t-tests were performed. Statistical significance is indicated as *p<0.05; **p<0.01; ***p<0.001; ****p<0.0001.

**Figure 3:**
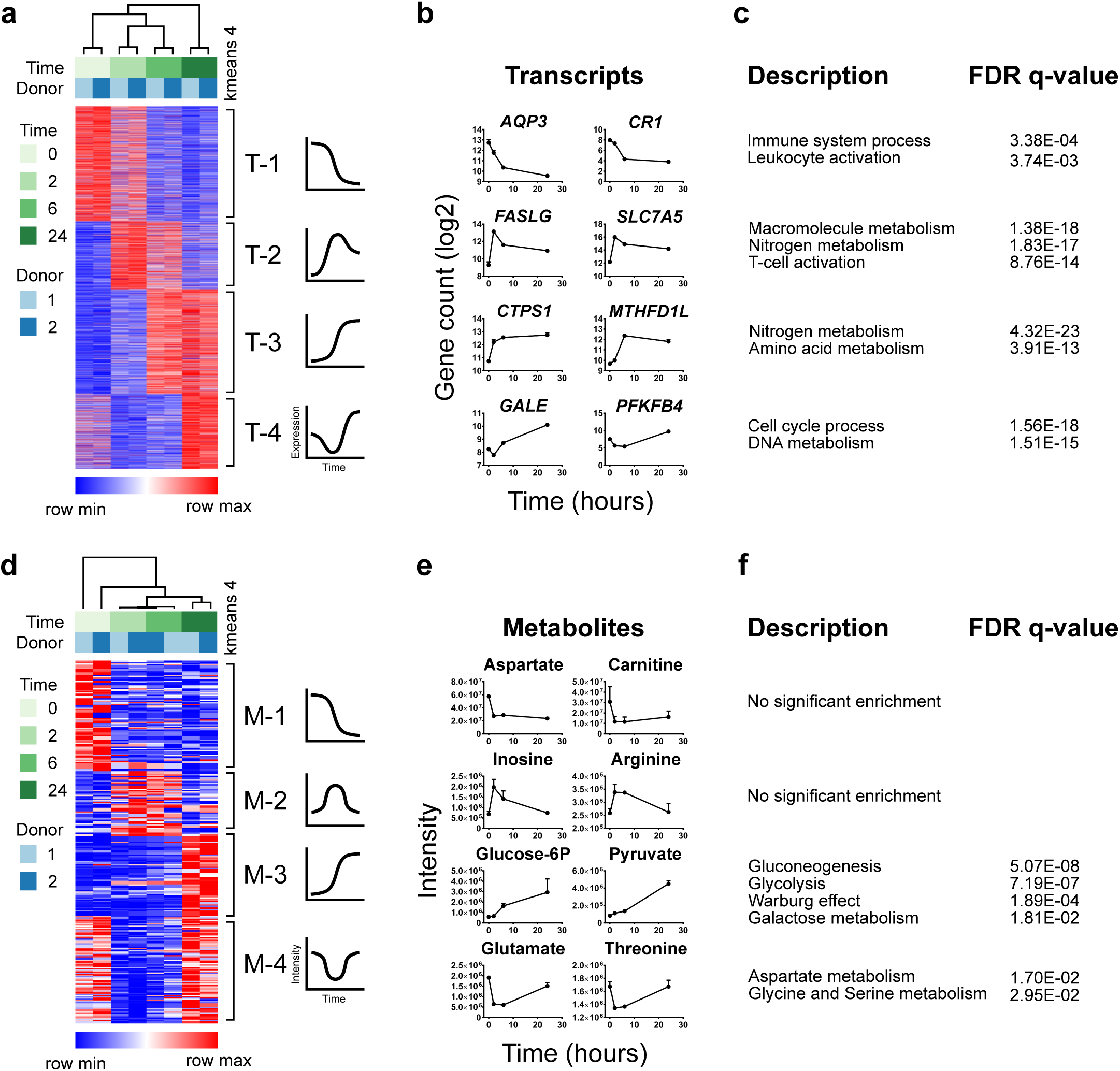
Kinetic transcriptomic and metabolomic profiling reveal expression of nitrogen and amino acid metabolism to be an early and sustained event upon TCR stimulation. Primary human CD4+ T cells from two independent donors were left rested (0) or were activated for 2, 6, or 24 hours with CD3/CD28 ImmunoCult. Cells were isolated for transcriptomic or metabolomic analysis. a, Heatmap of the top 3217 genes differentially expressed over time out of 7099 metabolically associated genes as defined by KEGG. Expression patterns of 4 K-means clusters (T-1 to T-4) are highlighted to the right. b, Expression of representative genes over time from each cluster. c, GO term enrichment (Description) and associated FDR q-value of each cluster with respect to the 7099 metabolically associated KEGG genes. d, Heatmap of the top 308 metabolites changing over time. Expression patterns of 4 K-means clusters (M-1 to M-4) are highlighted to the right. e, Expression of representative metabolites over time from each cluster. f, Over Representation Analysis (Description) and associated FDR q-value of each cluster with respect to Pathway-associated metabolite sets (MetaboAnalyst).

#### Transcriptomic data analysis

Transcriptional data was aligned using STAR^46^ (https://github.com/alexdobin/STAR) aligner. hg38 reference genome indexes for STAR aligner from GDC^48^ (https://api.gdc.cancer.gov/data/9c78cca3-c7d3-4cde-9d6c-54ec09421958) were used along with default settings of STAR aligner. Gene counts from resulting bam files were generated using featurecounts^47^ using GTF file from UCSC^49^ (http://bioinf.wehi.edu.au/featureCounts/). Differential expression between different cohorts was calculated using DESeq2^50^ package available with Bioconductor, only the top ten thousand transcripts were considered, sorted according to counts. DAVID^51^ was used to calculate the enrichment of different pathways in figure 3. GSVA^52^ package in R was used to calculate the pathway scores in figure 4a.

**Figure 4:**
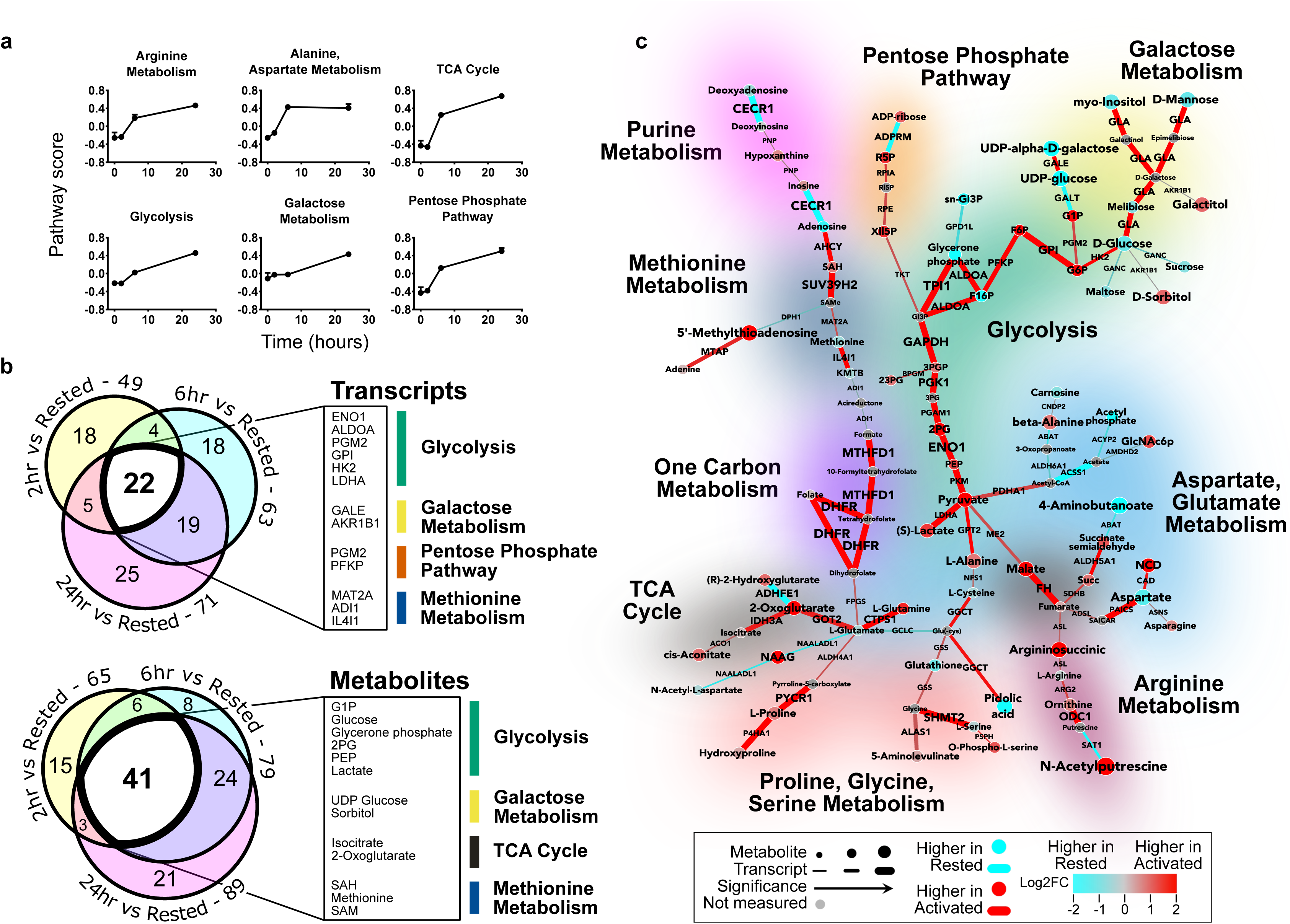
Integrated transcriptomic and metabolomic analysis reveal non-canonical reprogramming of metabolic pathways. Primary human CD4+ T cells from two independent donors were rested (0) or were activated for 2, 6, or 24 hours with CD3/CD28 ImmunoCult. Cells were isolated for transcriptomic or metabolomic analysis. a, GSVA indicating score of select pathways during activation time-course. b, Venn diagram of differentially expressed transcripts (top) or metabolites (bottom) at each activation time-point compared to rested cells. Specific transcripts or metabolites common to all activation time-points are highlighted with their respective pathway. c, Concordant metabolomics integrated with transcriptomics (CoMBI-T) analysis of 24 hour activation (red) vs rested cells (blue). Connecting lines represent transcriptional expression of enzymes, circular nodes represent metabolite expression, and increased width indicates higher significance. Specific pathways are highlighted.

#### Network Analysis (CoMBI-T)

Network analysis pipeline prescribed by Sergushichev et al.^53^ was used to construct the maps depicted in all figures. The network analysis pipeline suggested by Sergushichev et al. performs the integrated network analysis of transcriptional and metabolomic data to find the most changing subnetworks in the KEGG database. In order to compare CoMBI-T networks, the same FDR values were applied to the following figures: Supplementary figure 2, Supplementary figure 4, Figure 8. To compare two CoMBI-T networks and find rewiring of metabolism as in figure 5 we used Dynet^54^ analysis as devised by Goenawan et al.

**Figure 5:**
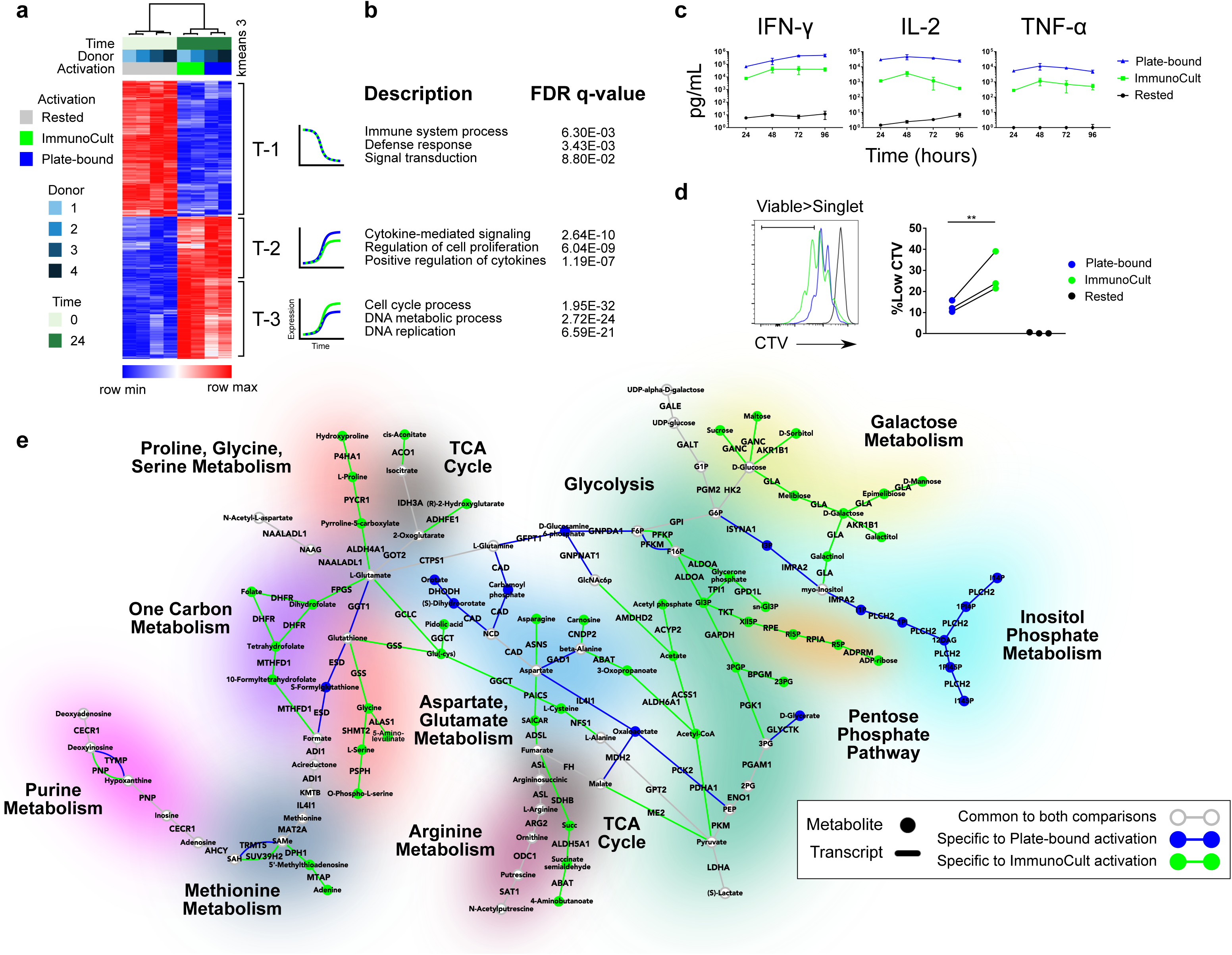
Strength of TCR stimulation differentially regulates metabolic reprogramming in human CD4+ T cells and correlates with differential pro-inflammatory phenotypes. Primary human CD4+ T cells from four independent donors were rested (0) or were activated for 24 hours with either CD3/CD28 ImmunoCult (green) or Plate-bound α-CD3/α-CD28 (blue). Cells were isolated for transcriptomic or metabolomic analysis. a, Heatmap of the top 1000 differentially expressed genes. Expression patterns of 3 K-means clusters (T-1 to T-3) are highlighted to the right. b, GO term enrichment (Description) and associated FDR q-value of each cluster with respect to all 21706 detected genes. c, Quantification of IFN-γ, IL-2, and TNF-α in supernatants collected at the indicated time points from one representative donor, n=3. d, CellTrace Violet staining of cells after 96 hours of activation from one representative donor (left). Cells were gated on viable singlets. Quantification of the CellTrace Violet-low population from three independent donors (right), each dot representing the average of one donor, n=3. Ratio paired two-tailed Student’s t-test was performed. Statistical significance is indicated as *p<0.05; **p<0.01; ***p<0.001; ****p<0.0001. e, DyNet comparison of CoMBI-T maps from 24 hour activation vs rested cells in either CD3/CD28 ImmunoCult or Plate-bound α-CD3/α-CD28. Connecting lines represent transcriptional expression of enzymes and circular nodes represent metabolite expression. Transcripts and metabolites common to both comparisons are highlighted in grey while those specific to either CD3/CD28 ImmunoCult or Plate-bound α-CD3/α-CD28 are highlighted in green or blue respectively. Specific pathways are highlighted.

#### Statistical Analysis

Data was analyzed using GraphPad Prism software. Data are presented as indicated in the figure legend which also highlights the statistical analysis used. Statistical significance is indicated in all figures by the following annotation: ns p≥0.05; *p<0.05; **p<0.01; **p<0.001; ****p<0.0001.

#### Data and Software Availability

All transcriptomic data is submitted to GEO with accession number GSE129829. All metabolomic data is submitted to metabolights^55^ with id MTBLS926. All data and CoMBI-T maps can be accessed online through an R shiny app available here: https://shashj199.shinyapps.io/shinyapp/

## Results

### Labeling analysis reveals increased utilization of glucose and altered metabolism of glutamine and fatty acids in activated human CD4+ T cells

Upon stimulation through their TCRs, human effector/memory CD4+ T cells undergo rapid cell division and concomitant production of inflammatory cytokines and chemokines. An increased metabolism is required to support these functions as is seen by the increased extracellular acidification rate (ECAR) and oxygen consumption rate (OCR) (Figure 1a-b). Previous studies showed that T cells require nutrients such as glucose and glutamine to fuel this increased metabolism^24^. To specifically investigate if glucose and glutamine were the predominant carbon sources required to fuel mitochondrial ATP production in human CD4+ effector T cells, we performed a mitochondrial fuel flexibility test (Figure 1c-d). These data demonstrated that pharmacological blockade of the mitochondrial pyruvate transporter (MPC) with UK5099 did not significantly suppress the OCR, a measure of mitochondrial respiration. These data might suggest that activated human T cells do not utilize glucose as the primary carbon source to fuel mitochondrial ATP production (Sup Figure 1a-b). Treatment with BPTES, an inhibitor for the enzyme glutaminase (GLS) required for glutamine entry into the TCA cycle, also failed to decrease the OCR (Sup Figure 1c-d). Etomoxir, an inhibitor carnitine palmitoyl transferase (CPT) required for mitochondrial uptake of long chain fatty acid derived carbons was the only pharmacological agent which reduced OCR of activated T cells. However, a recent report demonstrating a direct inhibitory effect of etomoxir on complex I of the electron transport chain reduces the utility of this agent in delineating the role of long chain fatty acids in T cell functions^25^. Activated T cells in the fuel flexibility test also exhibited a large residual OCR (∼100 pmol/min) despite the presence of all three inhibitors together (Figure 1c-d, Sup Figure 1). While the residual OCR might come from non-mitochondrial respiration, Sup Figure 7b suggests that non-mitochondrial respiration is minimal in activated T cells. These data led us to believe that the mitochondrial fuel flexibility test was not able to capture the dependence of the activated T cells on glucose and glutamine.

To determine the fate of glucose and glutamine, nutrients critical for effector functions of human T cells, we performed stable isotopic labeling in rested and acutely activated CD4+ T cells with either [U-^13^C_6_]-D-glucose or with [1,2-^13^C_2_]-L-glutamine. After introduction of [U-^13^C_6_]-D-glucose (Figure 2a), while the overall pools of pyruvate and lactate significantly increased upon activation, the ratio of labeled to unlabeled fractions were similar in rested and activated cells. Along with increased ECAR in activated T cells (Figure 1a), these data suggest a greater flux of glucose carbons through glycolysis upon activation without dilution of these pools through gluconeogenic or other contributing processes. The pool size of TCA cycle intermediates was also higher in activated T cells (Figure 2a). Overall, the percentage of labeled citrate was increased in activated cells compared to resting cells, but this increase in labeling was not consistently observed in other TCA intermediates such as malate suggesting increased dilution of the glucose-derived label (e.g., from glutamine or other unlabeled carbon sources).

In order to investigate these changing fractional enrichments, we used mass isotopomer multi-ordinate spectral analysis (MIMOSA) approach to perform steady state isotopomer ratio analysis on TCA cycle metabolites as described by Alves et al.^23^ Between pyruvate and glutamate (ϕPyrGlu) label was diluted by >80% in rested T cells and >90% in activated T cells (Sup Figure 2). This dilution indicates that unlabeled carbons enter somewhere between these two metabolites. In the absence of exogenous acetate, ketones, or branched chain amino acids, pyruvate dehydrogenase (PDH) and β-oxidation are the principal contributors to mitochondrial Acetyl-CoA (AcCoA). Under these conditions PDH will contribute labeled carbons while β-oxidation will contribute unlabeled carbons to AcCoA. In activated T cells we observed that PDH has a much greater contribution than β-oxidation to AcCoA indicated by a ϕPyrAc value of 0.8. In contrast, a ϕPyrAc value of 0.4 for rested T cells is a direct measure that the majority of AcCoA comes from unlabeled sources (i.e., beta-oxidation) (Figure 2b). Additionally, the value of ϕAcCit, which is indicative of the extent of dilution of citrate through unlabeled carbons from carboxylation of α-ketoglutarate (α-KG), shows a modest but significant decrease in activated T cells. The largest difference is observed in the value of ϕCitGlu, where the enrichment dropped 70% from citrate to glutamate in activated T cells compared to 25% in rested T cells. Since there are reversible reactions between citrate and glutamate, this additional dilution indicates that anaplerosis and/or exchange of glutamate with α-KG in activated cells was large relative to the flux within the TCA cycle compared to a much smaller exchange/anaplerosis in rested T cells. The estimated enrichment in oxaloacetate through pyruvate carboxylase (PC) was 2-3% in both states leading to a small ϕPyrOAA value indicating a negligible PC contribution to the TCA cycle in both rested and activated cells (Sup Figure 2).

In the presence of [1,2-^13^C_6_]-L-glutamine, there was a significant increase in the glutamine label in glutamate as well as TCA cycle metabolites in activated T cells (Figure 2d). The enrichments downstream from glutamine label also suggest a change in ATP-citrate lyase (ACLY) in activated T cells compared to resting cells as indicated by ϕCitMal (Figure 2e). The lack of enrichment in pyruvate and lactate (Figure 2d) also suggest negligible cataplerosis through oxaloacetate/malate by either PEPCK or Malic Enzyme (ME). Because the reversibility of the enzymes do not allow for quantitative measures, the increased steady state enrichment ratios (ϕGlnGlu, ϕGluCit) indicate a qualitative increase in glutamine exchange and/or anaplerosis.

Taken together, labeling and phenotypic data demonstrate an increased utilization of glucose in activated T cells. These data also reveal either increased anaplerosis or altered exchange reactions upon activation between glutamate and α-KG as well as altered fatty acid metabolism as observed through lowered dilution of AcCoA after glucose label and lowered dilution of malate after glutamine label.

### Integrated transcriptomic and metabolomic profiling reveals that TCR stimulation induces non-canonical reprogramming of metabolic pathways

To gain a mechanistic understanding of how TCR stimulation might augment anaplerosis, we profiled the kinetics of T cell transcriptome and metabolome changes. RNA-sequencing following 2, 6, or 24 hours of activation was performed from T cells from 4 human donors. Among 7000 metabolism-associated genes, we selected the top 3000 differentially expressed genes for K-means clustering (Figure 3a). Four distinct kinetic expression patterns were identified: decreased (T-1); transiently increased (T-2); stably increased (T-3); decreased at early time points and subsequently increased (T-4). Representative genes for each of the expression groups are shown in Figure 3b.

Gene ontology (GO) analysis of each group revealed several expected terms including immune cell activation (T-1, T-2, Figure 3c) and cell cycle (T-4, Figure 3c). Intriguingly, despite the profound metabolic changes, no canonical metabolic pathways were in the group T-1, where expression was decreased with TCR stimulation. Additionally, enrichment for genes of nitrogen and amino acid metabolism increased dramatically in T-2 and T-3 (Figure 3c) during the first few hours of activation and remained elevated. Non-targeted metabolomics were used to complement the transcriptional analysis where 308 metabolites changed upon TCR stimulation (Figure 3d). Four distinct metabolite clusters had kinetics mirroring the transcriptomic clusters in Figure 3a: M-1: metabolite abundance decreases over time; M-2: abundance peaks and returns to original levels; M-3: abundance increase over time; M-4: abundance dips early and increases over time (Figure 3e). Figure 3 provides a global picture of metabolism with limited focus on individual metabolites and genes.

The orthogonal transcriptomic and metabolomic data sets were integrated to reinforce influential metabolic pathways using CoMBI-T. Jha and colleagues have previously utilized to CoMBI-T to characterize global changes in metabolic pathways that accompany polarization of murine macrophages^26^. Geiger and colleagues utilized an analogous approach in which they integrated metabolomic and proteomic data sets to delineate the role of arginine metabolism in the regulation of human T cell survival^27^. Complementing the findings of these two reports, the data generated in this study describe rapid modulation of metabolic pathways in human T cells upon TCR stimulation and further identifies unexpected contribution of numerous amino acid metabolism pathways.

#### Kinetics of changes in metabolic processes upon TCR stimulation

We integrated the transcriptional and metabolomic data obtained at 2-, 6- and 24-hours post-stimulation (Figure 4a, Sup Figure 3) to identify metabolic genes whose transcription was modulated by TCR stimulation and to link them to changes in abundance of their known metabolite substrates and products in a pathway-agnostic approach. Through these analyses we observed significant changes in several metabolic modules. Many of these metabolic changes such as glycolysis, galactose metabolism, pentose phosphate pathway, TCA, Arginine metabolism, methionine metabolism, showed sustained upregulation at all time points as shown by GSVA analysis (Figure 4b-c).

#### Regulation of glucose metabolism upon TCR stimulation

Glycolysis appears as one of the most prominently changing metabolic modules in the CoMBI-T network, where expression of both upper and lower glycolytic genes is significantly upregulated (Sup Figure 4). Moreover, quantification of the extent of upregulation in glycolysis through GSVA reveals increased and sustained upregulation over time. CoMBI-T analysis also revealed metabolic modules of enzymes such as alpha-galactosidase (GLA) which promote conversion of glucose to other hexoses such as mannose, inositol, sorbitol, and galactitol (Figure 4c). The role of these hexoses in human CD4+ T cell function remain incompletely understood and warrants further investigation of these pathways.

#### Nucleotide synthesis

Consistent with the rapid proliferation that T cells are known to undergo, we observed changes in the expression of multiple enzymes and metabolites required for nucleotide synthesis (Figure 4c, upper and middle left). Several of these enzymes are known targets of clinical drugs such as methotrexate (dihydrofolate reductase; DHFR), or enzymes in which congenital mutations lead to clinical immunodeficiency such as adenosine deaminase 2 (CECR1) or CTP synthase 1 (CTPS1)^28^.

#### Amino acid metabolism

Consistent with results from our tracing studies we found several enzymes associated with amino acid metabolism to be strongly upregulated. The two overall themes suggested by the CoMBI-T analysis were a) biosynthesis of amino acids from both glucose and glutamine derived metabolites and b) the utilization of amino acids such as glutamine as substrates for anaplerotic filling of TCA cycle intermediates.

The first theme of biosynthesis of amino acids was readily exemplified through increased abundance of proline and alanine whose corresponding enzymes’ transcripts were also upregulated. These biosynthetic processes are well characterized and were mapped to be mediated by the canonical enzymes pyrroline-5-carboxylate reductase 1 (PYCR1) (Figure 4c, lower left) and alanine aminotransferase 2 (GPT2) (Figure 4c, center). The CoMBI-T algorithm also suggests that proline and alanine could possibly be derived from glutamate and pyruvate respectively, however this hypothesis needs to be tested in subsequent experiments.

A key exemplifier of the second theme was the increased expression of enzymes involved in glutamine utilization. Although the tracing studies pointed out that increase in labeled carbons in the TCA cycle coming from glutamine might be due to increased exchange or glutamine anaplerosis, there is a higher evidence for anaplerosis since transcripts of the enzymes responsible for glutamine to glutamate conversion (CTPS1 and GLS1) and glutamate to α-KG conversion (GOT2) are upregulated and appear in CoMBI-T. A key step in the utilization of glutamine is its deamination by GLS1. However, CoMBI-T analysis mapped the enzyme CTPS1 as a potential catalyst for glutamine to glutamate conversion because it is more significantly upregulated than GLS1 (although GLS1 is also strongly upregulated) (Figure 4c, lower left). The CTPS1 enzyme catalyzes CTP synthesis and during this process, transfers an amino group from glutamine to UTP to produce glutamate. While additional perturbational and tracing studies would be required to further substantiate the requirement for CTPS1 in the deamination of glutamine in activated human T cells, the results in this study suggest an efficient way for activated T cells to recycle nitrogen for nucleotide synthesis under metabolically demanding phases of cellular proliferation.

Two additional TCA intermediates that were increased in activated T cells were malate and fumarate. In agreement with the data reported by Geiger and colleagues^27^ we observed a strong accumulation of argininosuccinate, a key intermediate of the urea cycle. Notably, argininosuccinate seemed to be acting as a precursor of fumarate. Increased expression of the enzyme argininosuccinate lyase (ASL) supported a pathway for arginine utilization for accumulation of fumarate. Similarly, the concomitant upregulation of phosphoribosylaminoimidazole carboxylase (PAICS) and adenylosuccinate lyase (ADSL) suggested the conversion of arginine into fumarate (Figure 4c, lower right). Collectively, the data suggest that TCR-stimulation induces the expression of several novel metabolic nodes to synergistically promote two key features that have been described to accompany T cell activation; increased mitochondrial function and mass as well as nucleotide metabolism. These enzymes could be pharmacologically targeted for therapeutic modulation of the immune cells for inflammatory diseases. A list of the metabolic enzymes emphasized by the CoMBI-T analyses are provided in Supplementary Table 1.

### Strength of TCR stimulation differentially regulates metabolic reprogramming of human CD4+ T cells

Physiological stimulation of T cells is achieved in vivo through cognate interactions between the TCR expressed on mature T cells and a specific antigenic peptide presented in the context of MHC molecules expressed on the surface of antigen presenting cells. Mimicking these processes under *in vitro* cell culture conditions is quite challenging. High-throughput screening efforts required for drug discovery have conventionally used polyclonal stimuli that utilize stimulatory antibodies that cross link the TCR and costimulatory receptors such as CD28 to better simulate *in vivo* conditions in *in vitro* experiments. We compared the effects of two commonly used cross-linking formats which result in different strengths of TCR stimulation: 1) a weak TCR stimulation provided by the ImmunoCult reagent (α-CD3 and α-CD28 antibodies crosslinked with a soluble polymer) and 2) a stronger stimulation provided by well-established stimulation protocols in which α-CD3 and α-CD28 antibodies were immobilized on highly adsorbent tissue-culture plates (Plate-bound).

We first performed global transcriptomic analyses to compare the effects of these two stimulation conditions. Distinct gene expression patterns were obtained when the T cells were stimulated with either ImmunoCult or Plate-bound antibodies (Figure 5a). Whereas stimulation with Plate-bound antibodies gave a much stronger induction of genes enriched for the GO term ‘positive regulation of cytokine production’, cells stimulated with ImmunoCult exhibited increased expression of genes enriched for the GO term ‘DNA replication’ (Figure 5b). Analysis of cytokine production and cell proliferation confirmed these observations as stimulation with Plate-bound antibodies produced almost a log-fold increase in IFN-γ, IL-2 and TNF-α secretion compared to cells stimulated with ImmunoCult (Figure 5c). Conversely, cells stimulated with ImmunoCult exhibited increased proliferation compared to those stimulated with plate-bound antibodies (Figure 5d). CoMBI-T analyses revealed that the format used for TCR stimulation had distinct effects on metabolic reprogramming of human T cells. While there were numerous pathways that were similarly induced by ImmunoCult and Plate-bound formats of TCR stimulation, there were some notable differences (Figure 5e, Sup Figure 5). Expression of key enzymes required for inositol metabolism were found to be distinctly changed upon strong TCR stimulation provided by plate-bound antibodies. In contrast enzymes involved in amino acid metabolism were more strongly induced by the weaker stimulation provided by the ImmunoCult reagent. Furthermore, CoMBI-T analyses indicated that stimulation with the two different protocols resulted in different networks being adopted by T cells to facilitate similar metabolic transformations. A clear example could be visualized for one type of carbon metabolism (Figure 5e, middle left) involving the conversion of glutamate to formate. In the ImmunoCult activated cells, this conversion of glutamate to formate was facilitated through dihydrofolate reductase (DHFR), which formed the intermediary tetrahydrofolate, tetrahydrofolate synthase (MTHFD1) which subsequently catalyzed the conversion of tetrahydrofolate to formate. In contrast, upon Plate-bound activation glutamate was converted to glutathione by γ-glutamyltransferase 1 (GGT1), which was subsequently converted to formate by the activation of S-formylglutathione hydrolase (ESD). Collectively these data demonstrate that the nature and strength of TCR stimulation can have a differential impact on the regulation of T cell metabolism, although further studies are required to elucidate how closely these *in vitro* stimulation protocols mimic the metabolic changes that occur upon more physiologic stimulation of T cells *in vivo*.

### TCR-mediated increase in mTOR activity sustains metabolic reprogramming and pro-inflammatory functions of human CD4+ T cells

mTOR kinase is a kinase subcomponent of the mTORC1 complex that has been shown to promote protein translation and anabolic metabolism in numerous cell types. We (Sup Figure 6f) and others^29^ have observed a strong upregulation of mTOR activation upon stimulation of human T cells. As such, we investigated the contribution of mTOR kinase activation in the regulation of metabolism and function of activated human T cells.

Using the Seahorse cell analyzer, we measured the ECAR and OCR of resting and activated CD4+ T cells isolated from 9 independent human donors. The results demonstrate that shRNA-mediated inhibition of mTOR expression (Sup Figure 6) repressed activation-induced upregulation of aerobic glycolysis (and oxidative pentose phosphate pathway which also gives off acid) as well as mitochondrial respiration in T cells (Figure 6a-k, Sup Figure 7). As mTOR activity is very low in resting T cells, we saw no significant impact of mTOR knockdown on the respiration of resting T cells. Further dissection of the impact of mTOR knockdown revealed not only a suppression of ongoing glycolysis and mitochondrial respiration but also a strong impact on the maximal glycolytic capacity, maximal respiration, and ATP producing capability of activated CD4+ T cells (Figure 6a-k). These data suggested that mTOR activation is not only required for glucose uptake (as previously reviewed^30^) but is also important for expression of various enzymes required for glycolysis and mitochondrial respiration. Analysis of mitochondrial mass with MitoTracker green and levels of reactive oxygen species with CellROX dyes supported these observations and revealed an inability of CD4+ T cells transduced with mTOR shRNA to increase their mitochondrial mass and produce reactive oxygen species (Figure 6l-m).

**Figure 6:**
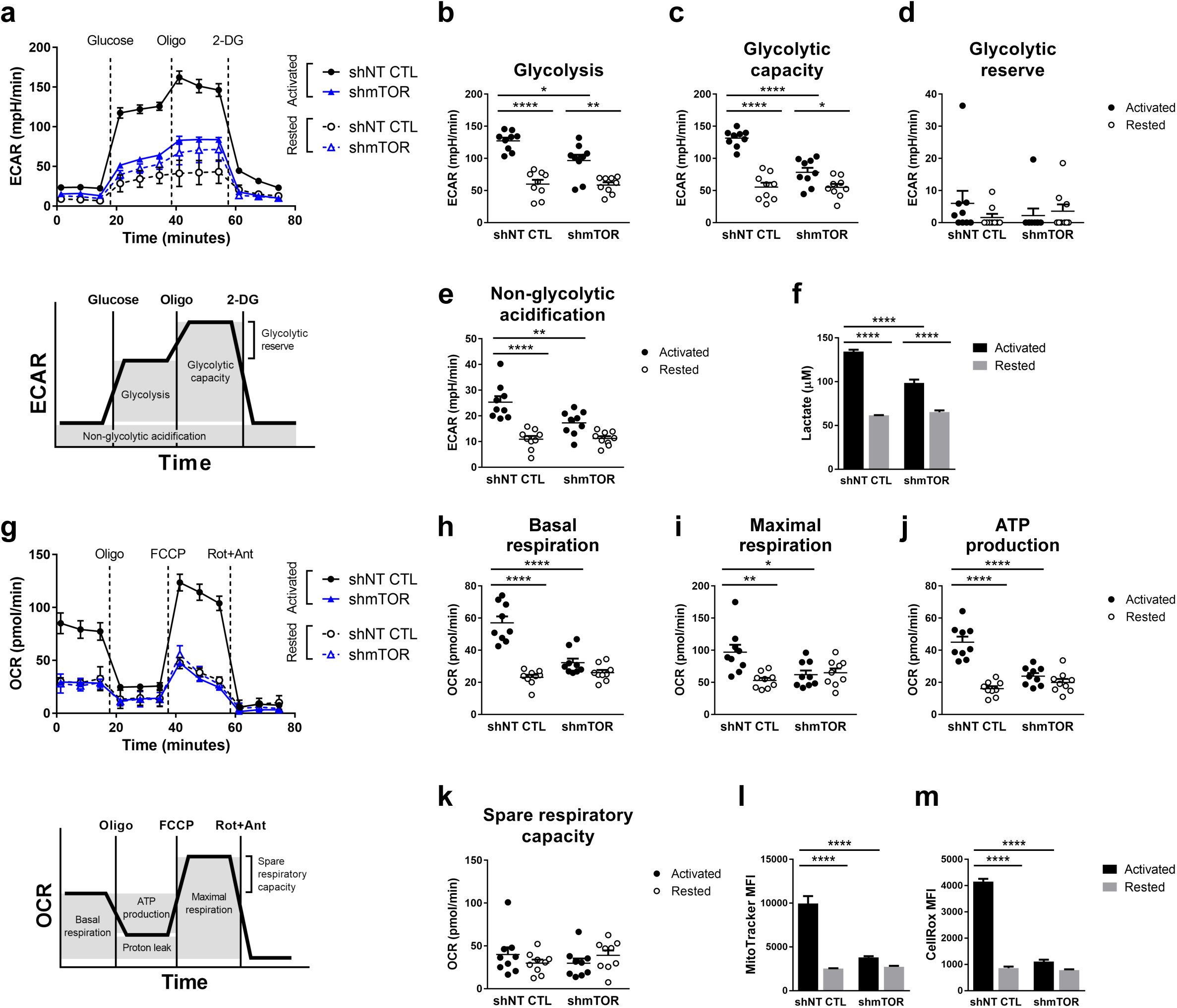
mTOR is required for increased glycolysis and mitochondrial respiration upon acute TCR activation. Primary human CD4+ T cells co-expressing ZsGreen and shNT CTL or shmTOR were rested or activated for 24 hours with CD3/CD28 ImmunoCult. a, ECAR measurements from one representative donor in response to glucose, oligomycin (Oligo), and 2-deoxyglucose (2-DG). Schematic below depicts the quantification of b, glycolysis, c, glycolytic capacity, d, glycolytic reserve, and e, non-glycolytic acidification from 9 independent donors, each dot representing the average from one donor, n=3. f, Quantification of lactate from one representative donor following the last ECAR measurement, n=3. g, OCR measurements from one representative donor in response to oligomycin, FCCP, and Rotenone+Antimycin (Rot+Ant). Schematic below depicts the quantification of h, basal respiration, i, maximal respiration, j, ATP production, and k, spare respiratory capacity from 9 independent donors, each dot representing the average from one donor, n=3. l, MitoTracker and m, CellRox staining from one representative donor, n=3. Cells were gated on ZsGreen-positive viable singlets. Data are presented as the mean ±SEM. Two-way ANOVA was performed with Tukey test to correct for multiple comparisons. Statistical significance is indicated as *p<0.05; **p<0.01; ***p<0.001; ****p<0.0001.

Due to such a broad impact of mTOR knockdown on T cell metabolism, we hypothesized that there would be a corresponding reduction in the pro-inflammatory functions of the CD4+ T cells. To test this hypothesis and confirm shRNA knockdown observations, we disrupted the mTOR gene in human CD4+ T cells using CRISPR-based reagents (Sup Figure 8). Intriguingly, while mTOR deletion did not lead to a complete block in pro-inflammatory functions of activated T cells, there was a significant and clear (< 80%) inhibition of cytokine production (Figure 7a-c). In contrast, deletion of ZAP70, a critical kinase required for early TCR signaling^31^, led to a complete block in both cytokine production and TCR-stimulated proliferation (Figure 7a-e, Sup Figure 9). We performed CoMBI-T analysis to determine if the decrease in pro-inflammatory functions we observed upon disruption of mTOR and ZAP70 expression were accompanied by attenuation of metabolic reprogramming. In keeping with a more pronounced impact of ZAP70 deletion on the pro-inflammatory functions of CD4+ T cells, ZAP70 deletion had a much stronger effect on expression of metabolic genes and abundance of their corresponding metabolites (Figure 7f-g). Collectively these data suggest that while mTOR activation is a key component of TCR-induced signaling cascades other TCR-induced signaling cascades also play non-redundant roles in rewiring of T cell metabolism.

**Figure 7:**
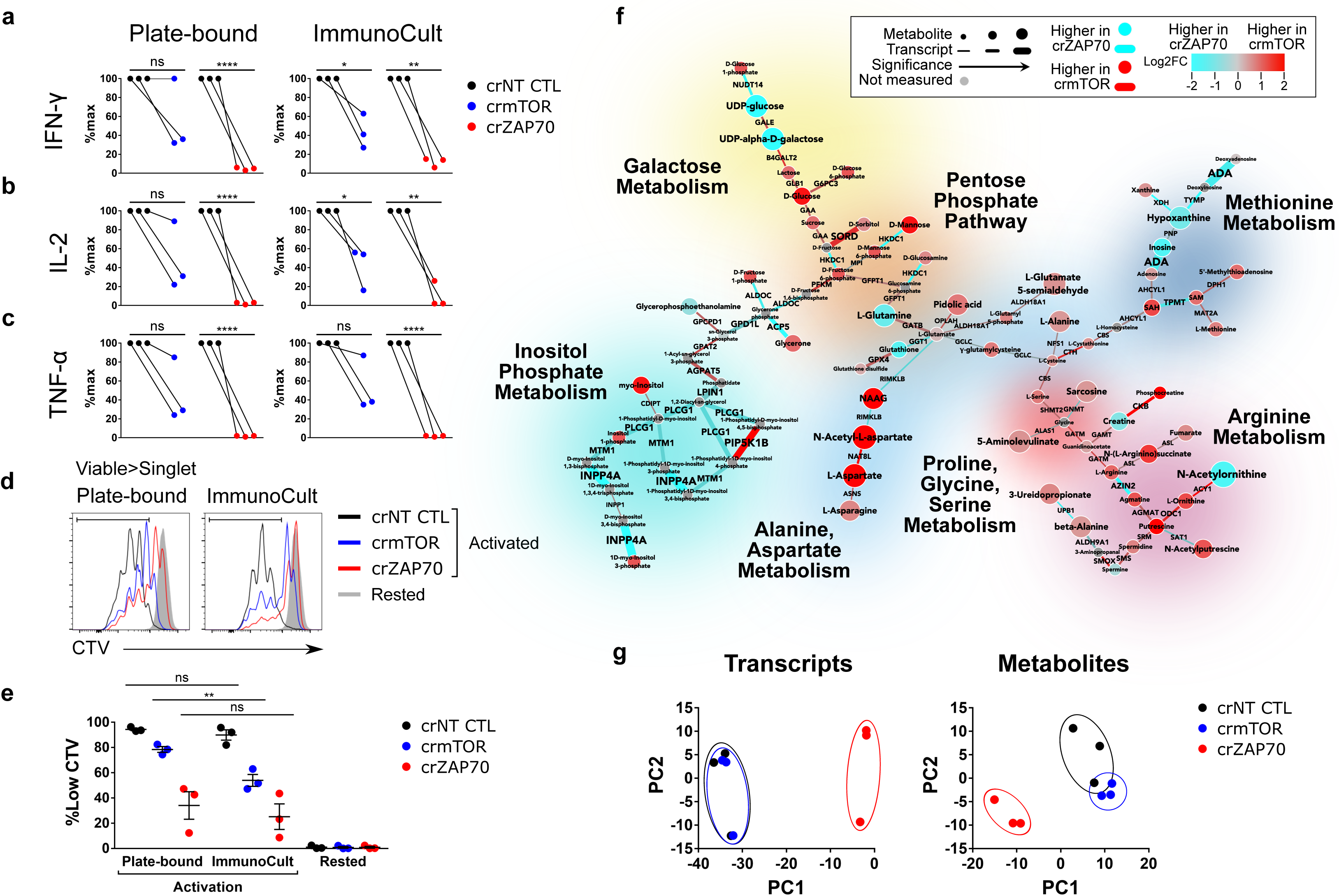
ZAP70 inhibition has a larger impact on T cell function and metabolism than mTOR inhibition. Primary human CD4+ T cells electroporated with CRISPR guides crNT CTL, crmTOR, or crZAP70 were rested or activated with either Plate-bound α-CD3/α-CD28 or CD3/CD28 ImmunoCult. Supernatants were collected after 24 hours for quantification of pro-inflammatory cytokines a, IFN-γ, b, IL-2, and c, TNF-α from three independent donors, each dot representing the average of one donor, n=3. d, CellTrace Violet staining of cells after 96 hours of activation from one representative donor. Cells were gated on viable singlets. e, Quantification of the CellTrace Violet-low population from three independent donors, each dot representing the average of one donor, n=3. Data are presented as the mean ±SEM. Paired two-tailed Student’s t-tests were performed. Statistical significance is indicated as *p<0.05; **p<0.01; ***p<0.001; ****p<0.0001. f, CoMBI-T analysis of crmTOR (red) vs crZAP70 (blue) after 24 hours of Plate-bound α-CD3/α-CD28 stimulation. Connecting lines represent transcriptional expression of enzymes, circular nodes represent metabolite expression, and increased width indicates higher significance. Specific pathways are highlighted. g, Principle component analysis of transcripts (left) or metabolites (right) after 24 hours of Plate-bound α-CD3/α-CD28 stimulation, each dot representing an independent donor.

### Targeting T cell metabolism enhances the immunosuppressive effects of anti-inflammatory drugs

The results of this study show that while ablating mTOR expression had a strong impact on the pro-inflammatory functions of T cells it was not sufficient to completely abrogate them. These data are consistent with studies that demonstrate that treatment with rapamycin (Rapa) in the context of allogeneic bone marrow transplantation delays pathogenic graft versus host disease rather than block it completely^32, 33^. Targeting ZAP70 for a more complete block of early TCR signaling has been challenging due to high levels of homology in the active site and ATP binding domains of these kinases, which poses difficulties in developing highly selective pharmacological inhibitors of ZAP70^34^. Therefore, we explored whether a combination of drugs such as cyclosporine (CsA) and Rapa which block distinct TCR-induced signaling pathways result in a more complete suppression of metabolism and inflammatory functions of human T cells.

We first evaluated the impact of each inhibitor alone on TCR-induced metabolic reprogramming using the CoMBI-T approach (Figure 8). In contrast to the results with mTOR deletion, we found that Rapa treatment had a modest impact on T cell metabolism. We observed inhibition of glutamine, glutamate. and arginine derived anaplerotic and biosynthetic processes, however we failed to see suppression of glycolytic genes (Figure 8a). These observations could be explained by the fact that the mTOR kinase is required for the function of both the mTORC1 as well as the mTORC2 complexes which have divergent and non-redundant anabolic functions. In contrast, Rapa has been demonstrated to completely block the function of the mTORC1 complex with a limited impact on the mTORC2 complex^35^.

**Figure 8:**
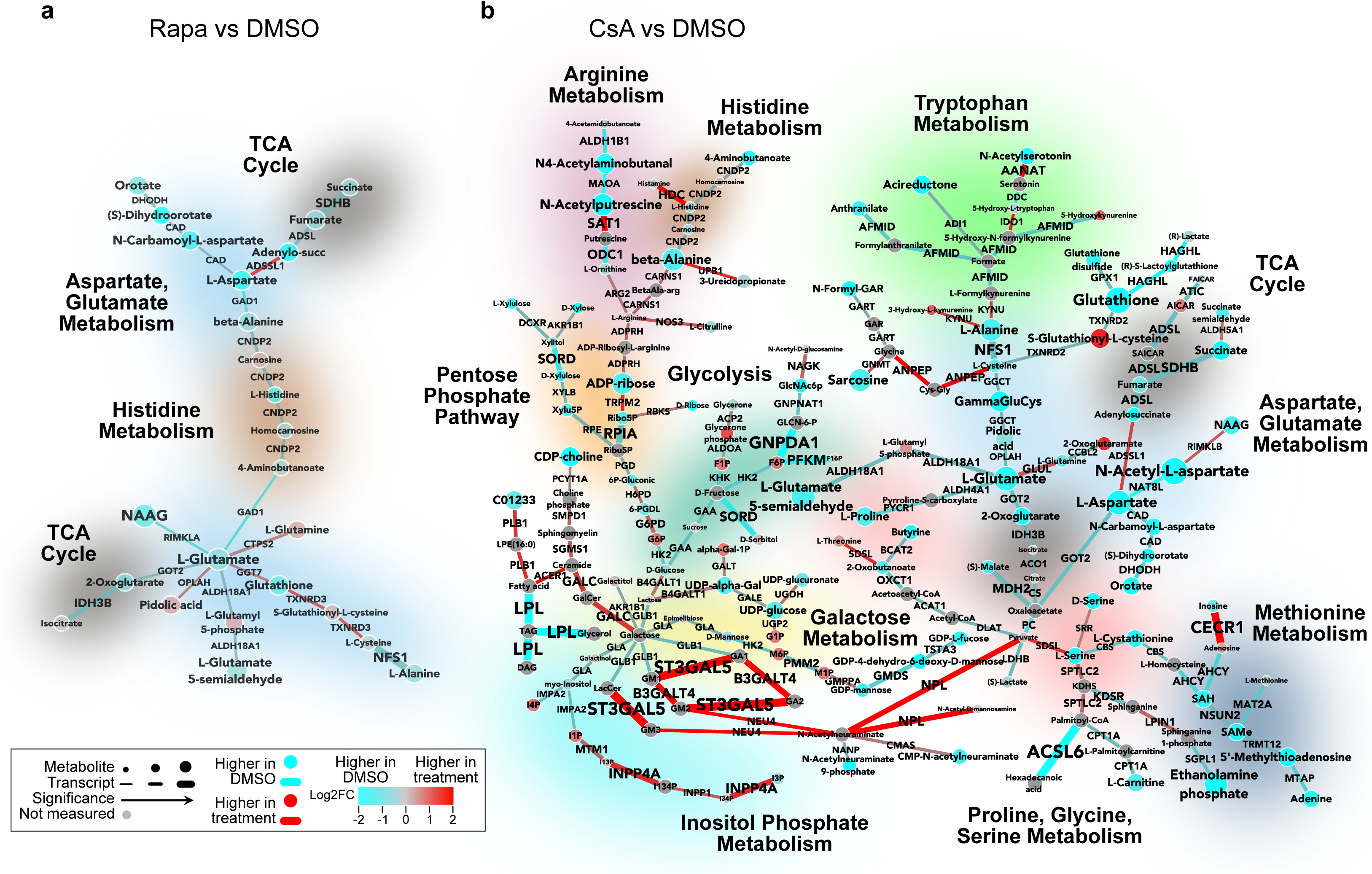
The anti-inflammatory drug Cyclosporine has a larger effect on T-cell metabolism than Rapamycin, a metabolic regulator. Primary human CD4+ T cells were activated for 24 hours with Plate-bound α-CD3/α-CD28 in the presence of vehicle (DMSO), Rapamycin (Rapa), or Cyclosporin (CsA). Cells were isolated for transcriptomic or metabolomic analysis. CoMBI-T analysis of a, Rapa (red) vs DMSO (blue) and b, CsA (red) vs DMSO (blue). Connecting lines represent transcriptional expression of enzymes, circular nodes represent metabolite expression, and increased width indicates higher significance. Specific pathways are highlighted.

Compared to Rapa, treatment with CsA alone had a much broader impact on the metabolic pathways in activated human CD4+ T cells suggesting that the upregulation of several metabolic enzymes was mediated through NFAT-driven transcription. Furthermore, we observed that the effects of CsA treatment were not limited to metabolic genes that we had previously found to be upregulated upon TCR stimulation. Specifically, we found that key genes associated with the pentose-phosphate pathway such as ribose-5-phosphate isomerase (RPIA), 6-phosphogluconate dehydrogenase (PGD) and Ribulose-phosphate 3-epimerase (RPE) were all strongly repressed by CsA treatment (Figure 8b). As these gene are not induced upon T cells activation, the data in this study suggest that the NFAT family of transcription factors are required for maintaining metabolic homeostasis in resting T cells as well.

We next evaluated how combined treatment of T cells with Rapa and CsA affects T cell metabolism and function. We activated human CD4+ T cells with ImmunoCult or Plate-bound antibodies in the presence of either CsA or Rapa alone or in combination. While CsA treatment was sufficient to completely block proliferation induced by weak ImmunoCult stimulation it was unable to halt proliferation induced by strong stimulation using Plate-bound antibodies. In keeping with earlier observations with our mTOR deletion experiments, Rapa treatment alone was also insufficient to completely block proliferation of activated T cells under both ImmunoCult and Plate-bound activation. However, combined treatment with Rapa and CsA suggested additive effects and led to a complete block in CD4+ T cell proliferation (Figure 9a-b). We further inquired if the additive effects of CsA and Rapa on T cell proliferation were reflected in the metabolic states. We performed CoMBI-T analysis of T cells which were stimulated with Plate-bound antibodies in the presence of either CsA or Rapa alone or in combination (Figure 9c). There was no evidence for any novel pathways being impacted with the combination. Rather, we observed that addition of Rapa further potentiated the immunosuppressive effects of CsA (Figure 9d). These data confirm observations that these inhibitors have an additive rather than synergistic effect on T cell metabolism and function. Collectively these data suggest that CoMBI-T analysis can be an efficient tool to determine the most appropriate combinations of drugs which can provide enhanced therapeutic modulation of immune responses while minimizing the impact on pathways whose inhibition could lead to increased adverse events and toxicity.

**Figure 9:**
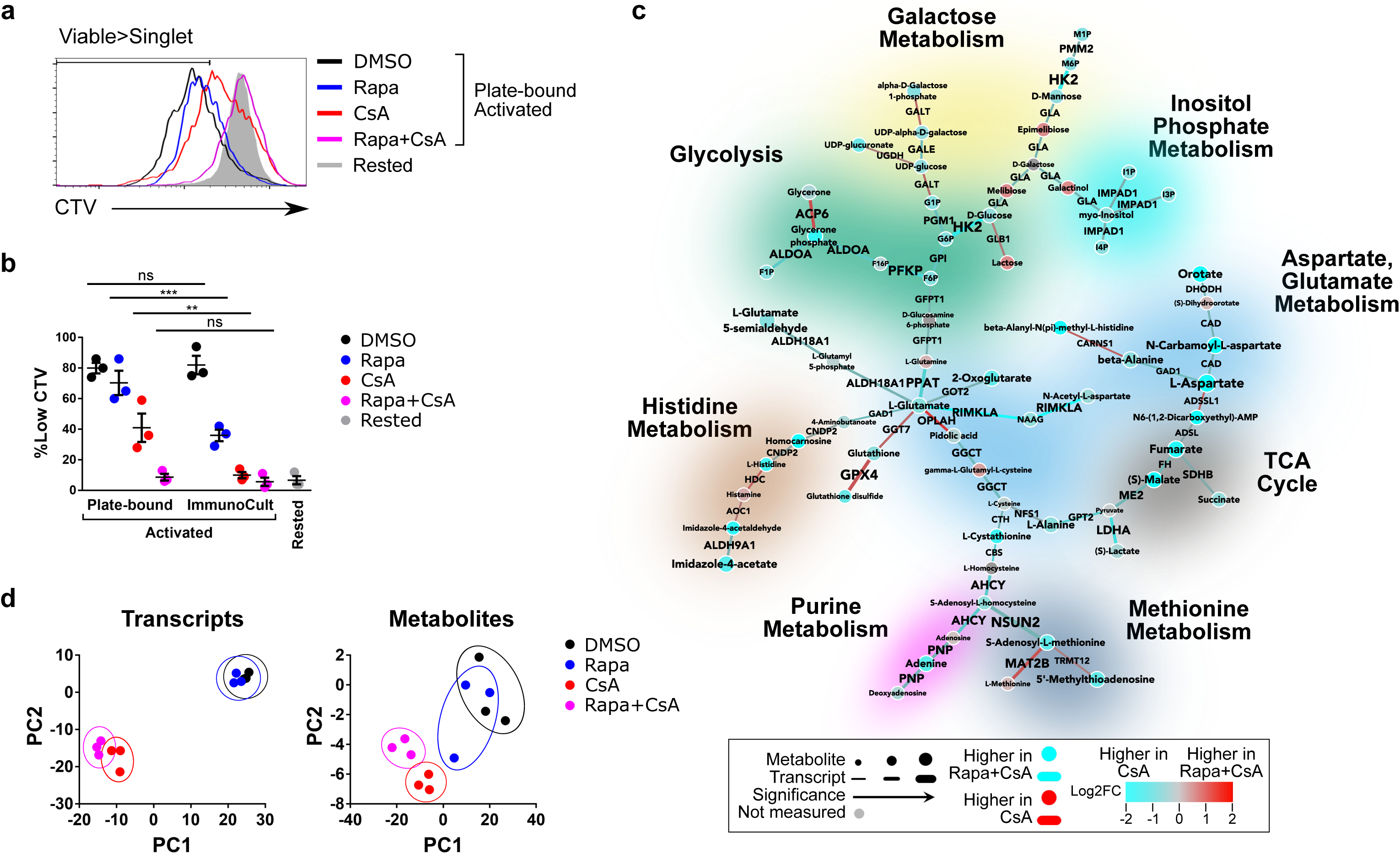
Targeting T cell metabolism using Rapamycin enhances the immunosuppressive effects of Cyclosporine. Primary human CD4+ T cells were rested or activated with either Plate-bound α-CD3/α-CD28 or CD3/CD28 ImmunoCult in the presence of vehicle (DMSO), Rapamycin (Rapa), Cyclosporin (CsA), or Rapa+CsA. a, CellTrace Violet staining of cells after 72 hours of activation from one representative donor. Cells were gated on viable singlets. b, Quantification of the Cell-Trace Violet-low population from three independent donors, each dot representing the average of one donor, n=3. Data are presented as the mean ±SEM. Paired two-tailed Student’s t-tests were performed. Statistical significance is indicated as *p<0.05; **p<0.01; ***p<0.001; ****p<0.0001. c, CoMBI-T analysis of CsA (blue) vs Rapa+CsA (red) after 24 hours of Plate-bound α-CD3/α-CD28 stimulation. Connecting lines represent transcriptional expression of enzymes, circular nodes represent metabolite expression, and increased width indicates higher significance. Specific pathways are highlighted. d, Principle component analysis of transcripts (left) or metabolites (right) after 24 hours of Plate-bound α-CD3/α-CD28 stimulation, each dot representing an independent donor.

## Discussion

In this study we investigated the contribution of two key nutrients, glucose and glutamine, to the pro-inflammatory functions of human effector T cells. Pharmacological blockade of either the mitochondrial pyruvate transporter (MPC) and/or glutaminase (GLS1) did not suppress the increase in OCR following activation of human effector CD4+ T cells. Combinatorial addition of these inhibitors further precluded the possibility that human effector T cells were highly flexible in their fuel choice for glucose and glutamine and could circumvent the requirement of either of these nutrients by solely relying on the other.

In contrast, OCR of activated T cells was reduced by etomoxir, an inhibitor of carnitine palmitoyl transferase (CPT1a), an enzyme to be transported into the mitochondria. While, it is tempting to speculate from these data that long chain fatty acids are the primary fuel source for human effector T cells, recent data have demonstrated that etomoxir may not be specific and may also inhibit complex 1 of the mitochondrial electron transport chain^25^. Further experiments would be required to more specifically establish the requirement for fatty acids for mitochondrial respiration.

It is possible that either the pharmacologic blockade could be incomplete or that GLS and MPC may be dispensable for mitochondrial ATP production. The former is supported by the carbon-tracing experiments that clearly demonstrate that both glucose-as well as glutamine-derived carbons can enter into the metabolites of the TCA cycle. MIMOSA and CoMBI-T analyses suggest that deamination of glutamine is increased to generate glutamate in activated T cells which could indeed fuel mitochondrial ATP production. Since 80% of the mitochondrial AcCoA comes from pyruvate, the combination oxidation and anaplerosis from both glucose and glutamine, but not beta-oxidation, appear to be dominant as T cells become activated.

Transcriptomic and metabolomic profiling studies complement the above data and demonstrate that upon T cell activation several novel pathways are induced to synergistically promote nucleotide synthesis and TCA anaplerosis. In principle, these pathways could compensate for the inhibitory effects of BPTES on glutaminase activity in activated T cells. The importance of one such enzyme, CTPS1, is underscored by clinical presentation of impaired T cell immunity in patients with congenital CTPS1 deficiency^28^. These data suggest that modulation of metabolic pathways could alter immune cell function and provide a rationale for therapeutic inhibition of key metabolic enzymes for ameliorating inflammatory diseases.

CoMBI-T analyses reveal several pathways such as the glutamine metabolism, urea cycle, and polyol metabolism which are specifically upregulated upon T cell activation. Inhibitors of a key component of the polyol pathways, aldo-keto reductase (AKR1B1) which were initially developed for the treatment of diabetic nephropathy^36^ have been demonstrated to have anti-inflammatory activities in pre-clinical models of inflammation^37–40^. Additionally, Geiger and colleagues have demonstrated that arginine metabolism in activated T cells is critical for their survival^27^. Finally, results from this study, which suggest that glutamine is a key precursor for biosynthetic reactions that not only promote TCA anaplerosis but also DNA and proline synthesis, could provide a rationale for the efficacy of DON, a glutamine inhibitor in ameliorating rejection of allogeneic transplants in mice as well as protecting mice from inflammation associated-encephalomyelitis induced by alphavirus infection^41, 42^. Collectively, these data suggest that the CoMBI-T analyses described here point toward several new enzymes which could be targeted for inflammatory diseases. A list of these candidate enzymes is provided in Supplementary Table 1.

The knockdown and deletion experiments performed in this study confirmed that among the numerous pathways triggered upon TCR stimulation, activation of mTOR kinase is a key pathway which drives the metabolic reprogramming required for the pro-inflammatory functions of human effector T cells. In agreement with previous reports^43^, we observed a critical role of mTOR not only in the upregulation of glycolytic pathway genes but also in the upregulation of genes involved in fatty acid metabolism (Sup figure 10). CoMBI-T analyses further revealed additional pathways that required mTOR activity. Specifically, we found that upregulation of ornithine aminotransferase (OAT) and ornithine decarboxylase (ODC1), enzymes required for polyamine metabolism, was impaired in the absence of mTOR activity. Similarly, expression of glutamate and proline biosynthesis as well as TCA pathway genes were attenuated correlating with both decreased ECAR and OCR of activated T cells transduced with mTOR shRNA vectors. Further studies would be required to determine if these pathways are critical for the function of T cells or contribute to adverse events such as oral mucositis and hypertriglyceridemia observed upon clinical use of mTOR inhibitors.

While the effect of mTOR inhibition on T cell metabolism was broad, it was insufficient to completely block the proinflammatory functions of T cells. This study demonstrated that ZAP70 deletion attenuated expression of additional metabolic pathways which were distinct from those affected by mTOR deletion. These data suggest that additional mTOR-independent signaling cascades triggered upon TCR stimulation also play important roles in metabolic reprogramming of activated human T cells.

Rapamycin (Rapa) and cyclosporine (CsA) are two clinically utilized immunosuppressants which are often used in combination for the prevention and treatment of transplant associated GVHD. Using CoMBI-T analysis we evaluated the impact of these agents on T cell metabolism. Contrary to expectations, we found a broader impact of CsA treatment on T cell metabolism. Part of this could be due to the fact that Rapa affects only one of the two complexes, mTORC1, which require mTOR kinase activity. Intriguingly, we also found that CsA impacts the expression of numerous metabolic genes which are expressed at steady state levels in T cells and not further increased upon T cell stimulation. Combinatorial treatment with both inhibitors did not impact any additional metabolic pathways but did increase the extent to which these pathways were suppressed, supporting an additive rather than synergistic effect of combining these two drugs.

We propose that CoMBI-T analyses could inform the development of novel combinatorial regimens which could synergistically target immune cell metabolism to suppress their inflammatory functions without dose-limiting adverse events.

---

**Sup Figure 1:**
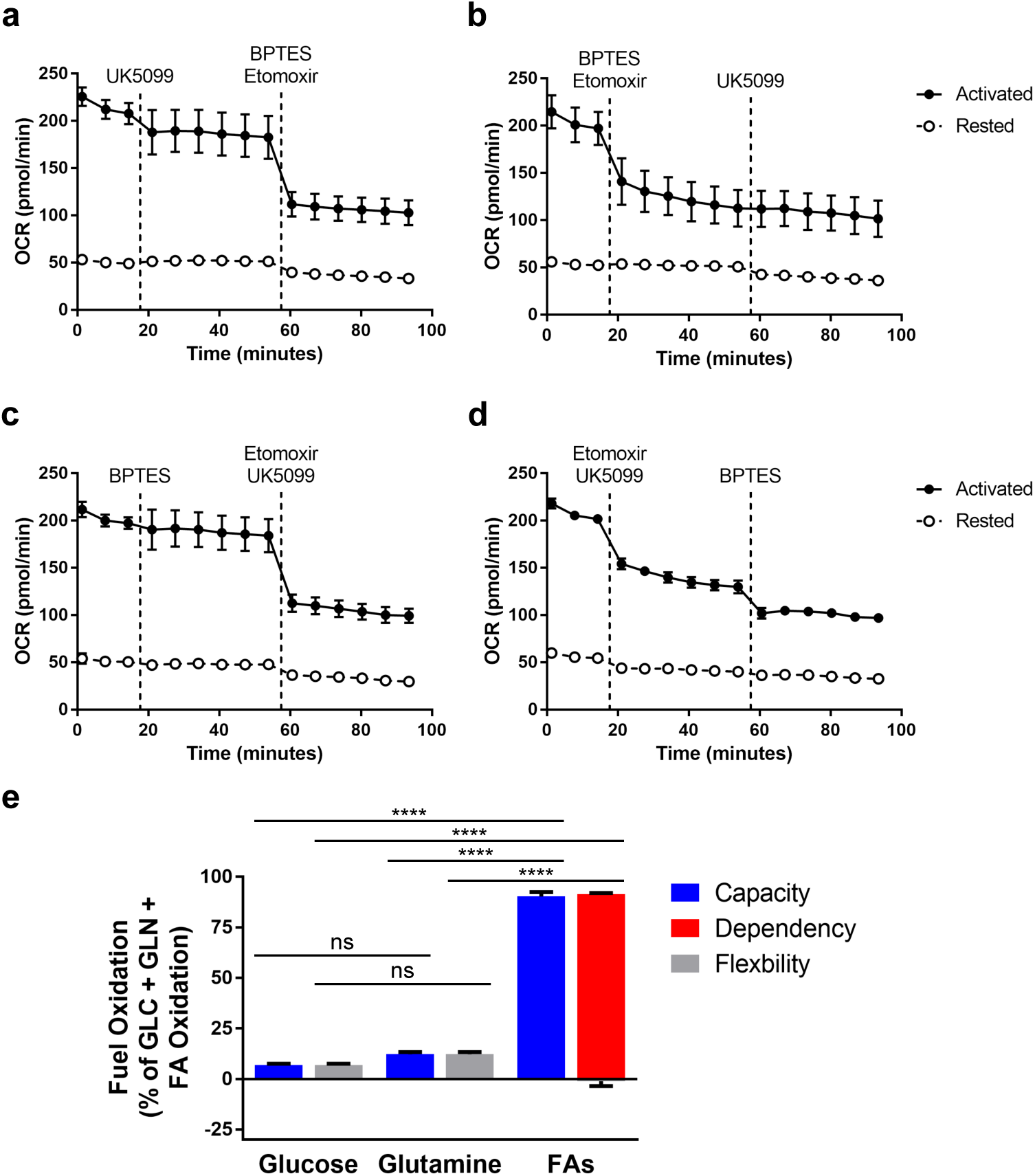
Utilization of glucose through MPC or glutamine through GLS1 are dispensable for mitochondrial ATP production in activated human CD4+ T cells. Primary human CD4+ T cells were rested or activated for 24 hours with CD3/CD28 ImmunoCult. OCR measurements from one representative donor in response to UK5099, BPTES, and Etomoxir indicating glucose a, dependency and b, capacity; and glutamine c, dependency and d, capacity. e, Quantification of glucose, glutamine, and fatty acid capacity, dependency, and flexibility in an additional donor. Data are presented as the mean ±SEM, n=3. Two-way ANOVA was performed with Tukey test to correct for multiple comparisons. Statistical significance is indicated as *p<0.05; **p<0.01; ***p<0.001; ****p<0.0001.

**Sup Figure 2:**
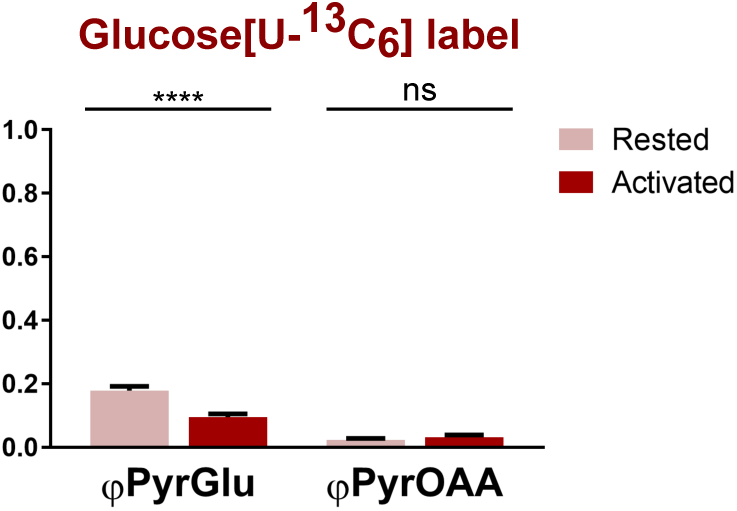
Glucose-derived carbon is diluted out of TCA cycle between the metabolites pyruvate and glutamate. Primary human CD4+ T cells from one donor were rested or activated for 24 hours with CD3/CD28 ImmunoCult. Cells were labeled with [U-^13^C_6_]-D-glucose. Phi values of select metabolites are illustrated. Data are presented as the mean ±SD, n=6. Unpaired two-tailed Student’s t-tests were performed. Statistical significance is indicated as *p<0.05; **p<0.01; ***p<0.001; ****p<0.0001.

**Sup Figure 3:**
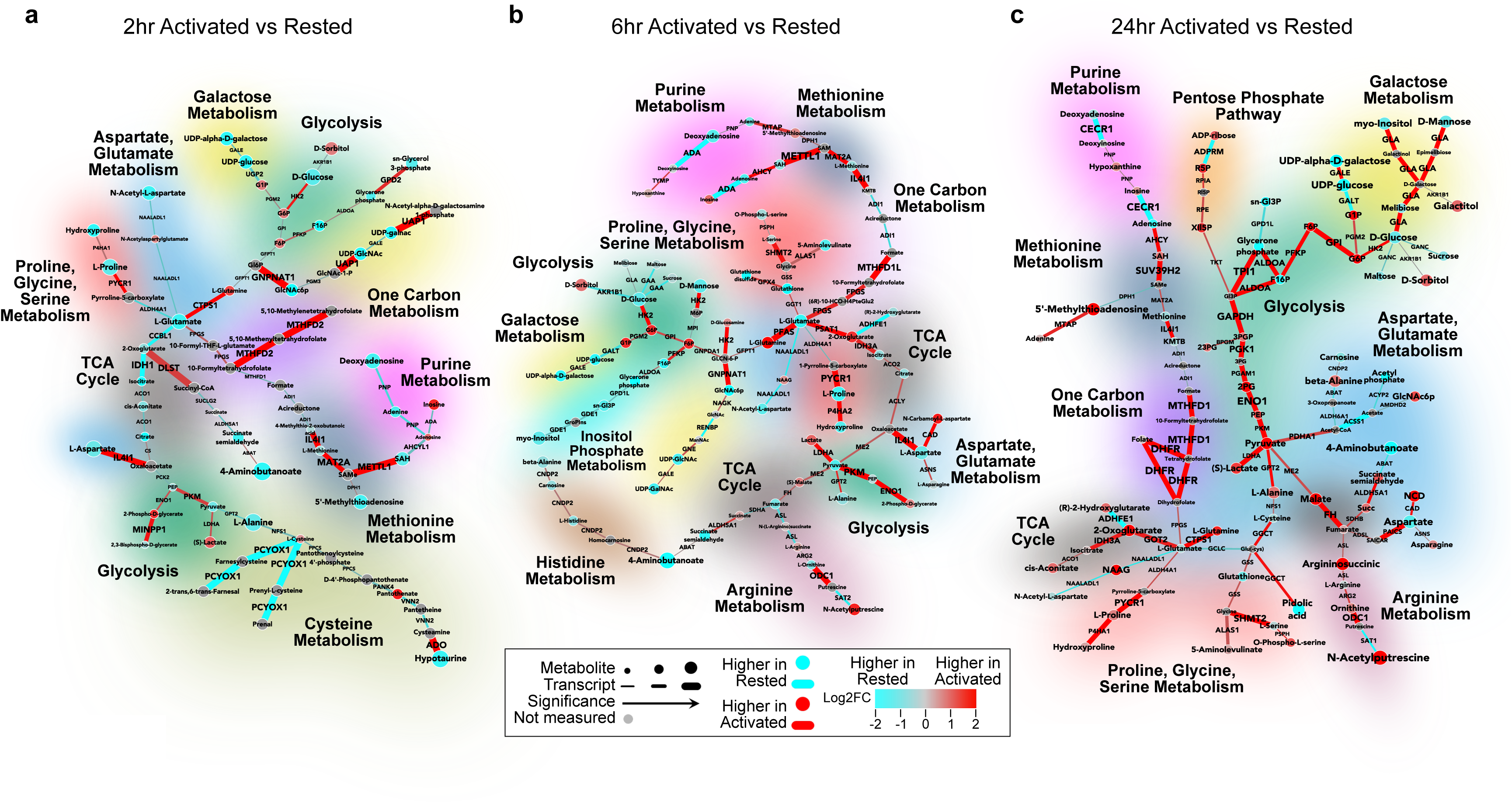
Kinetic profiling of integrated transcriptomic and metabolomic analysis reveal early and sustained changes in metabolic pathways. Primary human CD4+ T cells from two independent donors were rested (0) or were activated for 2, 6, or 24 hours with CD3/CD28 ImmunoCult. Cells were isolated for transcriptomic or metabolomic analysis. CoMBI-T analysis of a, 2 hour, b, 6 hour, or c, 24 hour activated cells (red) vs rested cells (blue). Connecting lines represent transcriptional expression of enzymes, circular nodes represent metabolite expression, and increased width indicates higher significance. Specific pathways are highlighted.

**Sup Figure 4:**
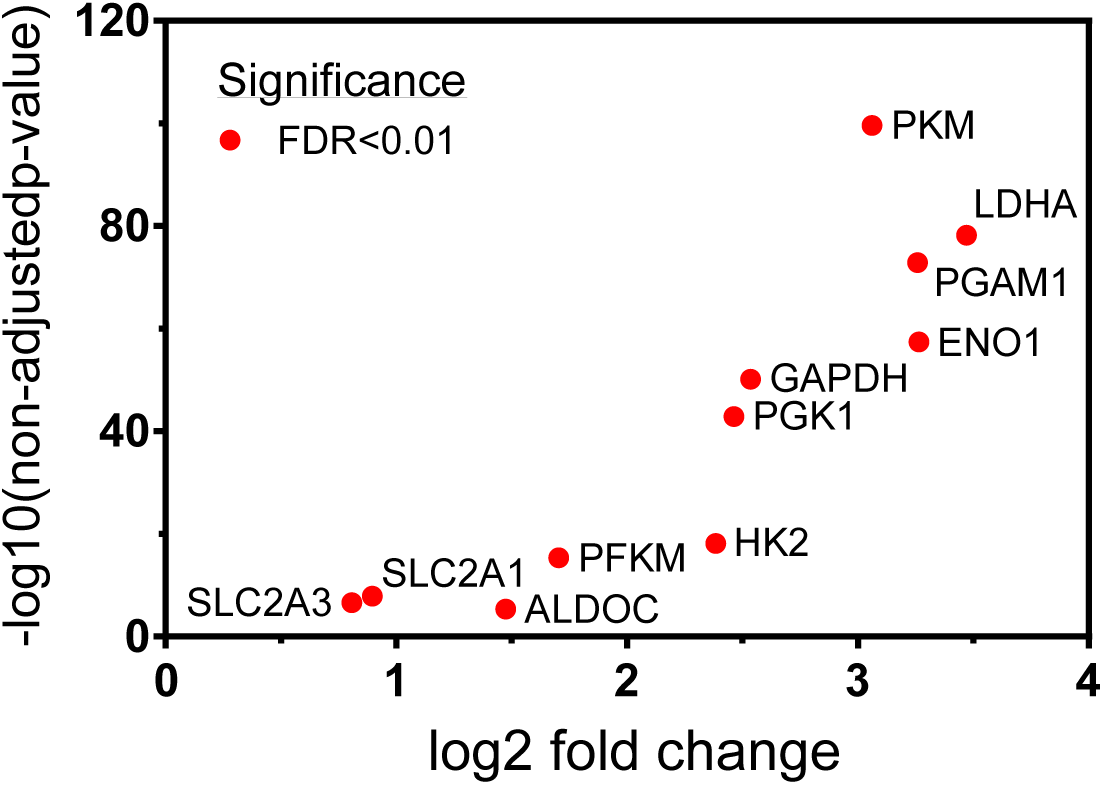
Expression of glycolytic enzymes upon TCR stimulation. Primary human CD4+ T cells from two independent donors were left rested (0) or were activated for 24 hours with CD3/CD28 ImmunoCult. Cells were isolated for transcriptomic analysis. Log 2-Fold Change plotted for select glycolytic enzymes after 24 hours of activation compared to rested cells.

**Sup Figure 5:**
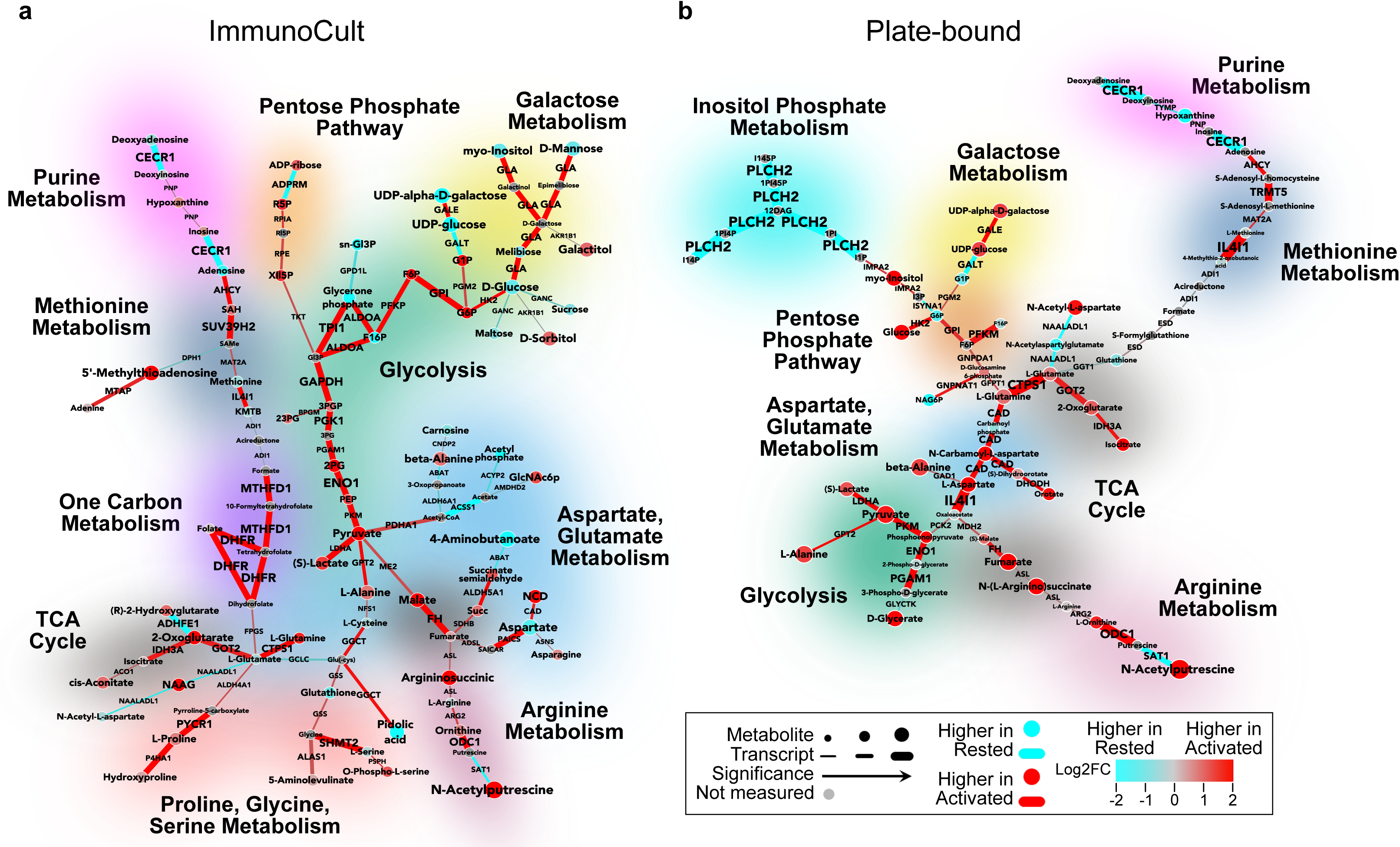
Strength of TCR stimulation differentially regulates metabolic reprogramming in human CD4+ T cells and correlates with differential pro-inflammatory phenotypes. Primary human CD4+ T cells from two independent donors were rested (0) or were activated for 24 hours. Cells were isolated for transcriptomic or metabolomic analysis. CoMBI-T analysis of a, CD3/CD28 ImmunoCult, or b, Plate-bound α-CD3/α-CD28 activated cells (red) vs rested cells (blue). Connecting lines represent transcriptional expression of enzymes, circular nodes represent metabolite expression, and increased width indicates higher significance. Specific pathways are highlighted.

**Sup Figure 6:**
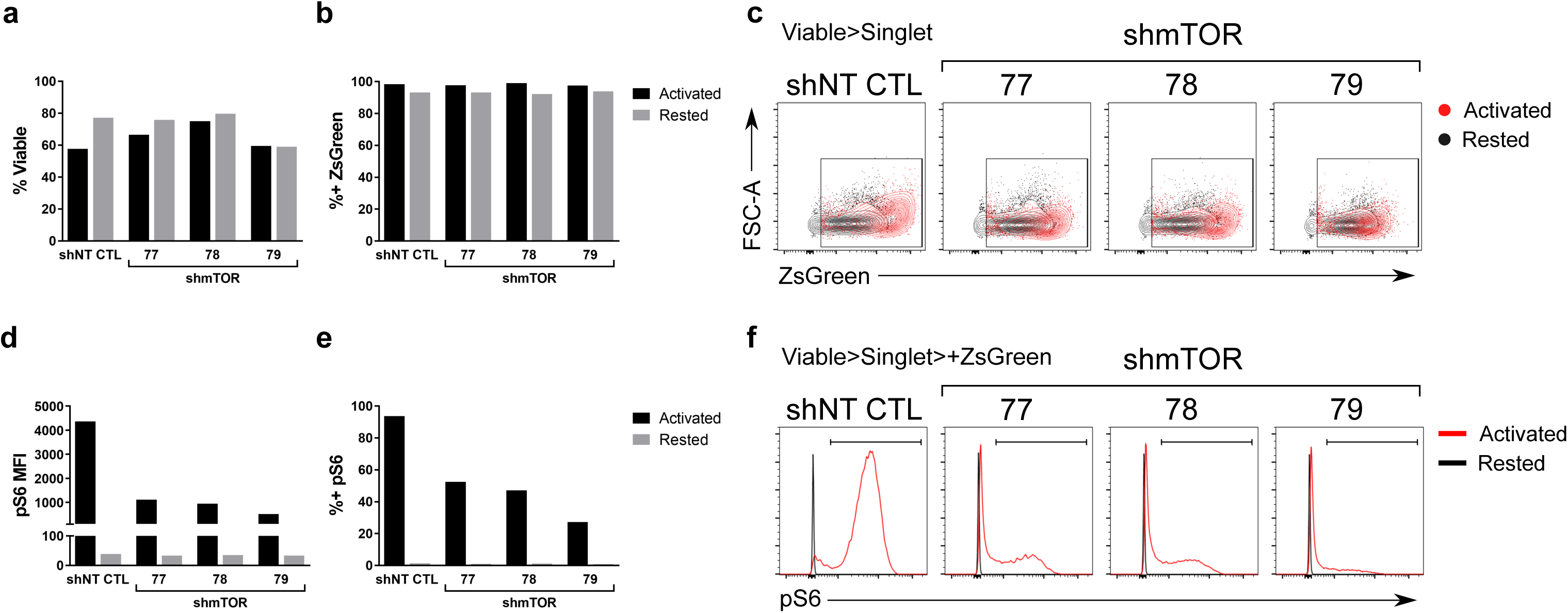
Efficiency of lentiviral-mediated shRNA knockdown in primary human CD4+ T cells. Primary human CD4+ T cells co-expressing ZsGreen and shNT CTL or one of three shmTOR hairpins were rested or activated for 24 hours with CD3/CD28 ImmunoCult. a, Viability of transduced cells after acute activation. b, Percent of cells expressing ZsGreen, gated on viable singlets. c, Dot plots of ZsGreen, gated on viable singlets. Phospho-S6 expression represented as d, MFI and e, percent positive cells, gated on ZsGreen-positive viable singlets. f, Representative histograms of phospho-S6 staining.

**Sup Figure 7:**
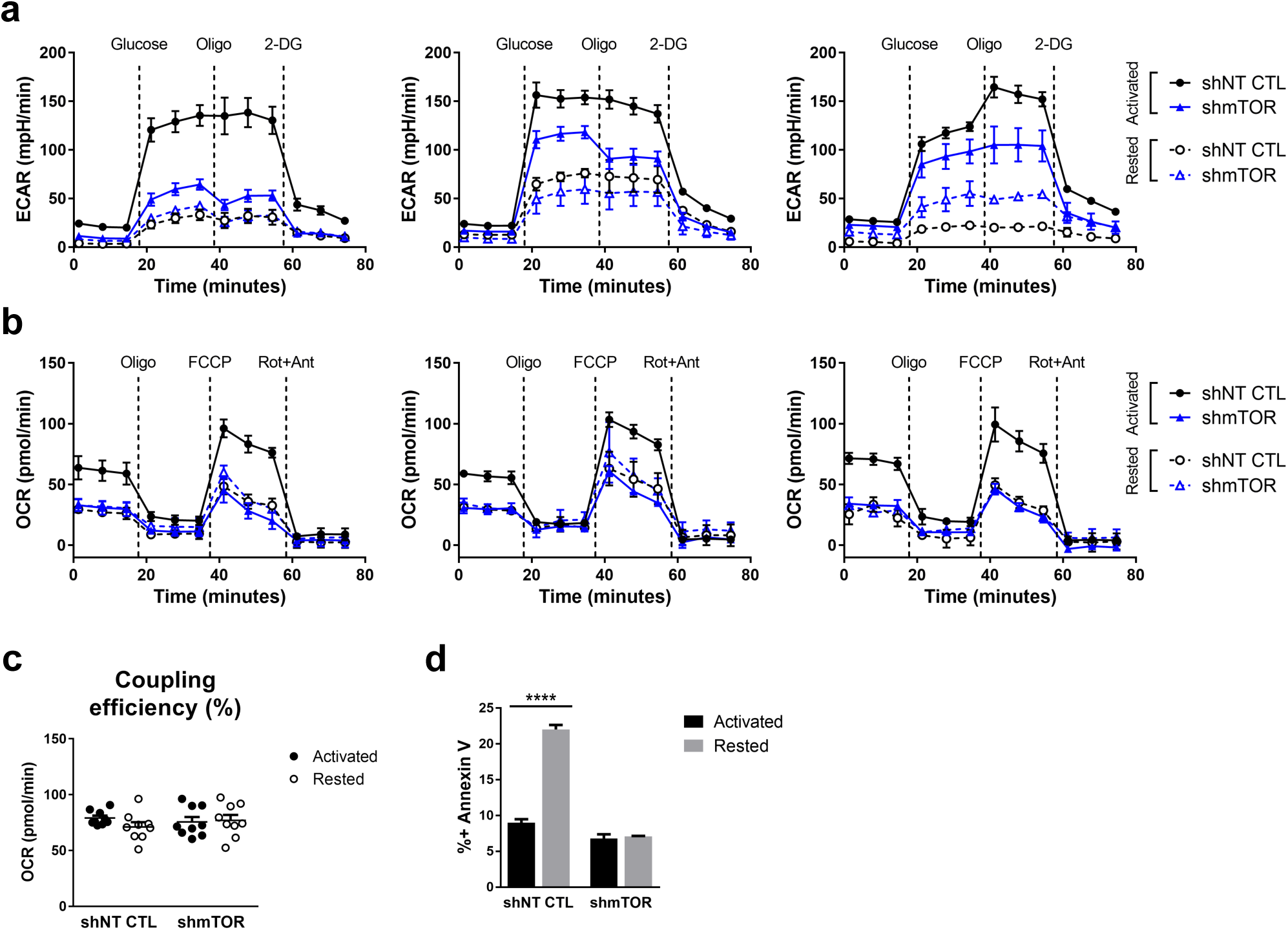
mTOR is required for increased glycolysis and mitochondrial respiration upon acute TCR activation. Primary human CD4+ T cells co-expressing ZsGreen and shNT CTL or shmTOR were rested or activated for 24 hours with CD3/CD28 ImmunoCult. a, ECAR measurements from three additional donors in response to glucose, oligomycin (Oligo), and 2-deoxyglucose (2-DG). b, OCR measurements from three additional donors in response to oligomycin, FCCP, and Rotenone+Antimycin (Rot+Ant). c, Quantification of coupling efficiency. d, Annexin V staining from one representative donor, n=3. Two-way ANOVA was performed with Tukey test to correct for multiple comparisons. Statistical significance is indicated as *p<0.05; **p<0.01; ***p<0.001; ****p<0.0001.

**Sup Figure 8:**
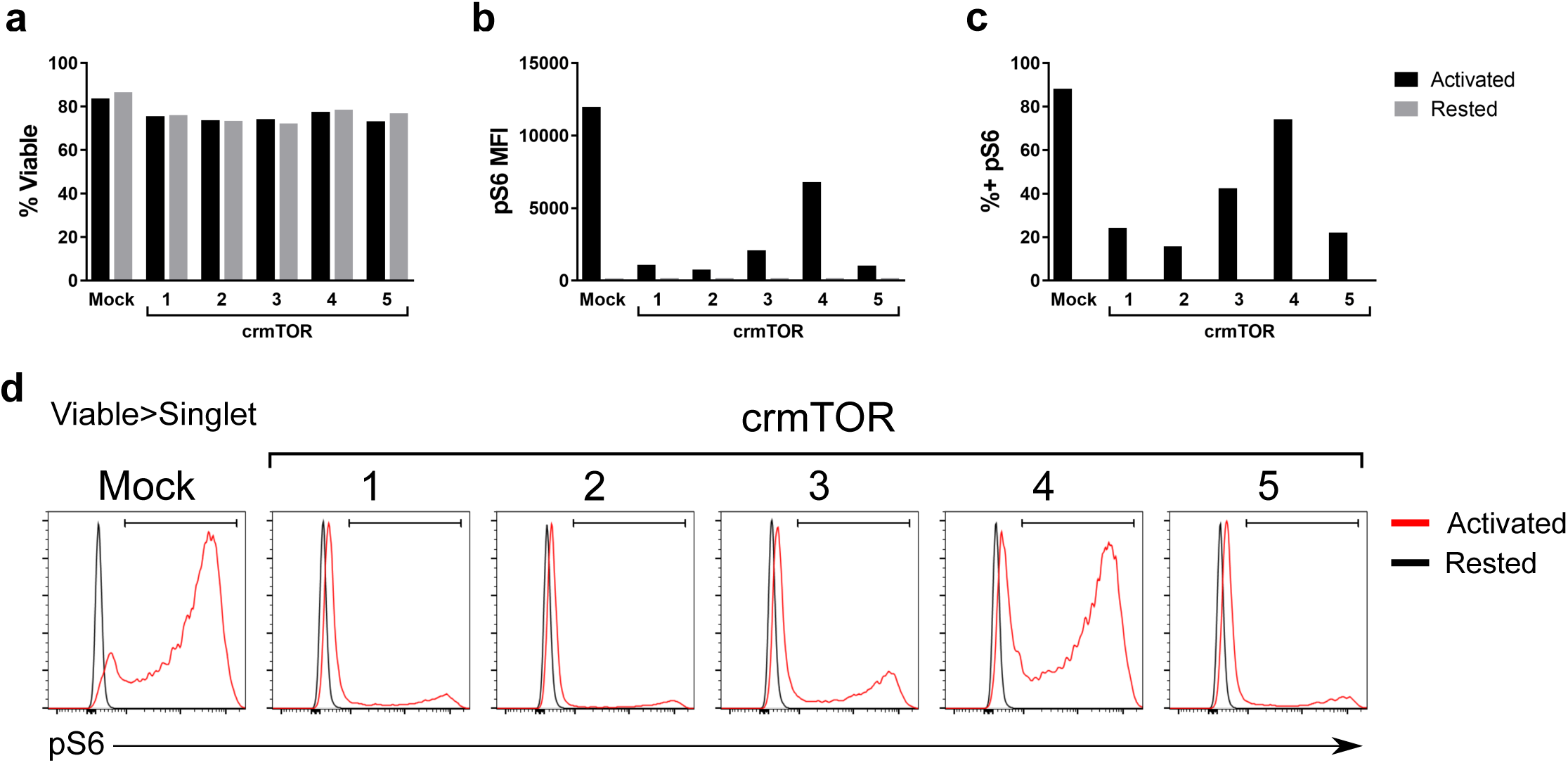
Efficiency of CRISPR-Cas9 mediated knockout of mTOR in primary human CD4+ T cells. Primary human CD4+ T cells electroporated with one of five crmTOR CRISPR guides or mock were rested or activated for 24 hours with CD3/CD28 ImmunoCult. a, Viability of electroporated cells after acute activation. Phospho-S6 expression represented as b, MFI and c, percent positive cells, gated on viable singlets. d, Representative histograms of phospho-S6 staining.

**Sup Figure 9:**
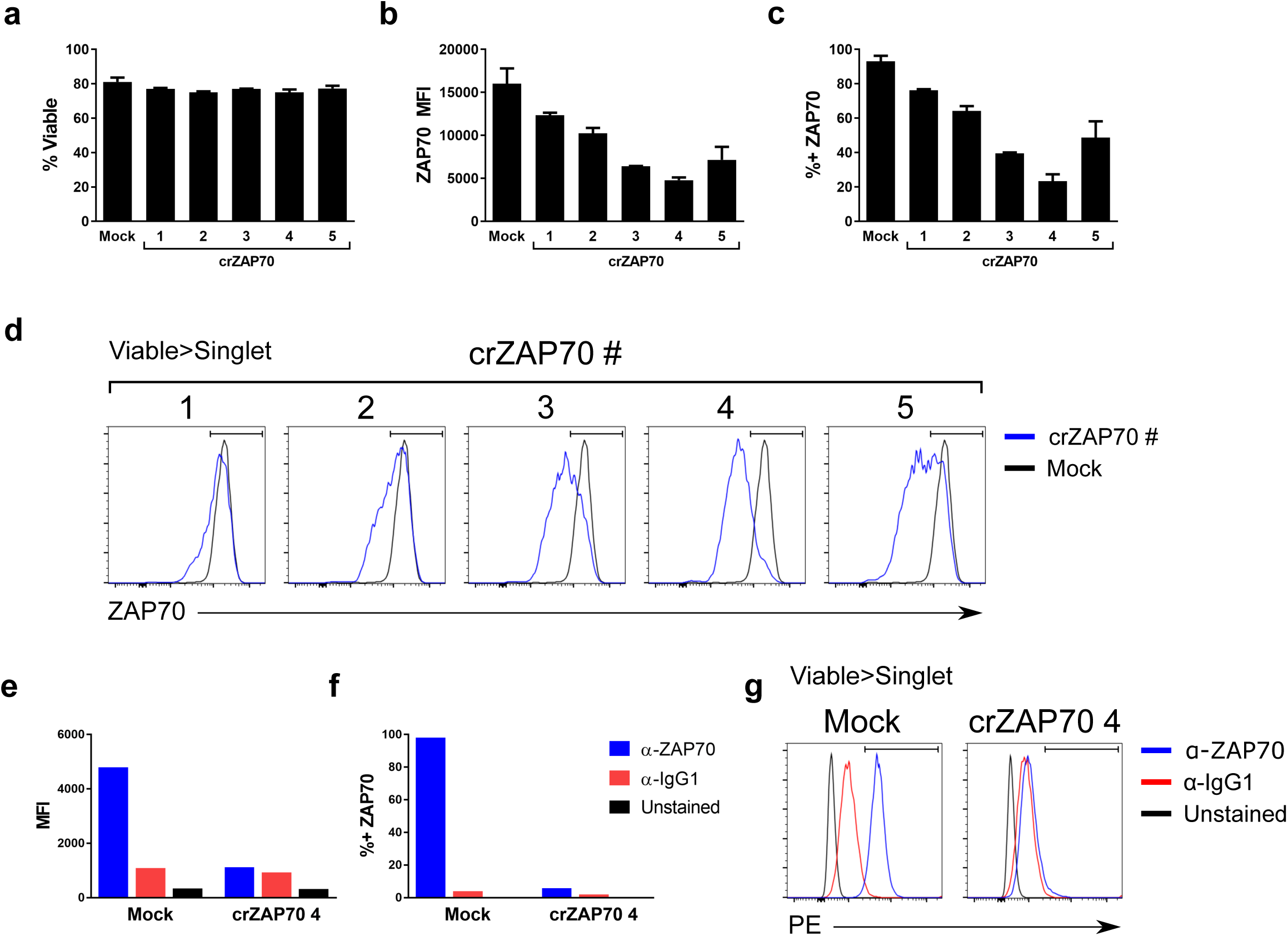
Efficiency of CRISPR-Cas9 mediated knockout of ZAP70 in primary human CD4+ T cells. Primary human CD4+ T cells were electroporated with one of five crZAP70 CRISPR guides or mock. a, Viability of electroporated cells. ZAP70 expression represented as b, MFI and c, percent positive cells, gated on viable singlets. d, Representative histograms of ZAP70 staining. e, MFI and f, percent positive cells, gated on viable singlets, after staining with ZAP70 antibody or isotype control in mock or crZAP70 #4 knockout cells. g, Representative histograms of ZAP70 and isotype control staining.

**Sup Figure 10:**
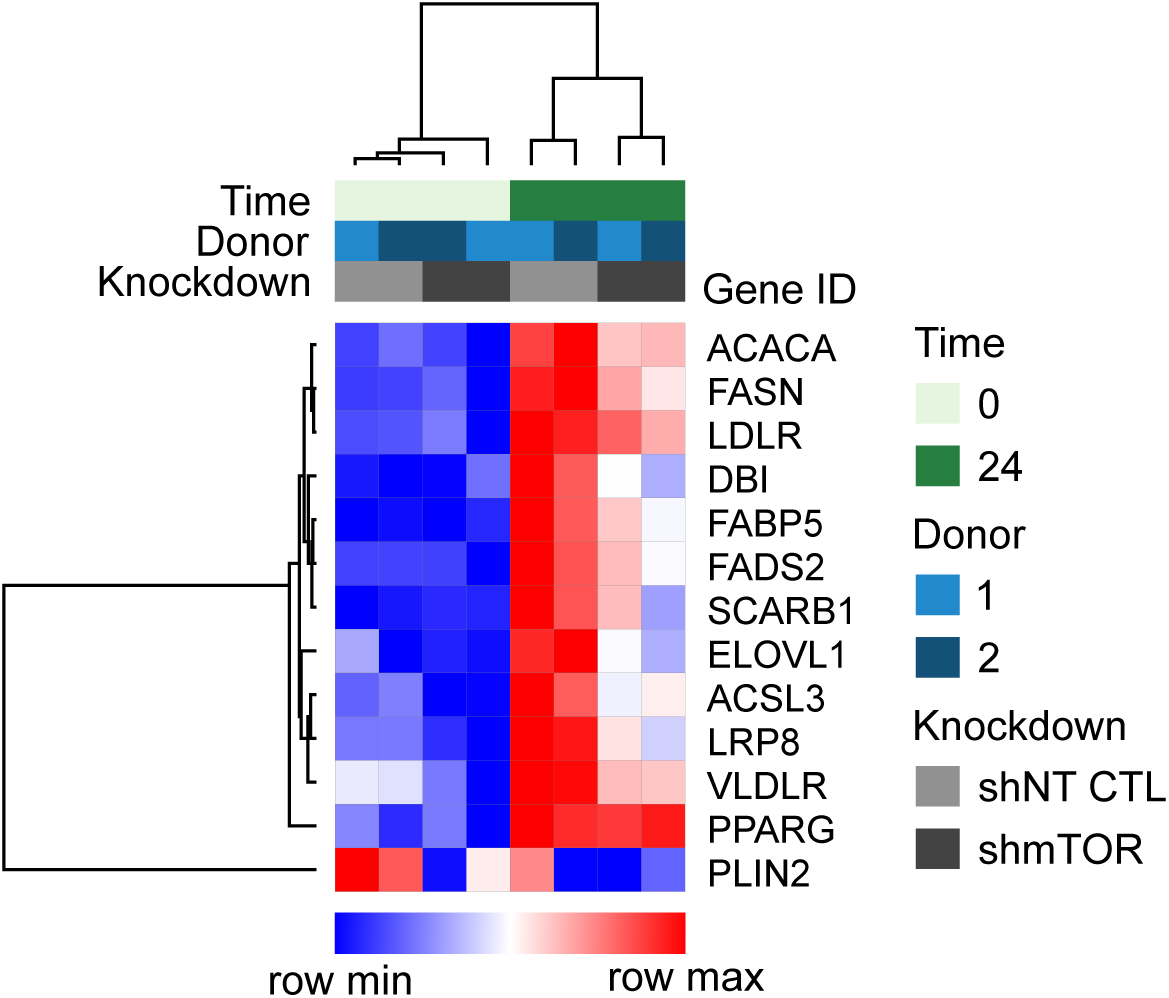
mTOR is required for upregulation of genes related to fatty acid metabolism upon acute TCR activation. Primary human CD4+ T cells from two independent donors co-expressing ZsGreen and shNT CTL or shmTOR were rested or activated for 24 hours with CD3/CD28 ImmunoCult. Cells were isolated for transcriptomic analysis.

## Notes

#### Summary of Updates

Correcting Figure 9 legend.

https://shashj199.shinyapps.io/shinyapp/

https://www.ncbi.nlm.nih.gov/geo/query/acc.cgi?acc=GSE129829

https://www.ebi.ac.uk/metabolights/MTBLS926

## References

1. Lee, Y. S., Wollam, J. & Olefsky, J. M. An Integrated View of Immunometabolism. Cell 172, 22–40 (2018).

2. Wang, R. et al. The Transcription Factor Myc Controls Metabolic Reprogramming upon T Lymphocyte Activation. Immunity 35, 871–882 (2011).

3. Jacobs, S. R. et al. Glucose Uptake Is Limiting in T Cell Activation and Requires CD28-Mediated Akt-Dependent and Independent Pathways. J. Immunol. 180, 4476–4486 (2008).

4. Cao, Y., Rathmell, J. C. & Macintyre, A. N. Metabolic Reprogramming towards Aerobic Glycolysis Correlates with Greater Proliferative Ability and Resistance to Metabolic Inhibition in CD8 versus CD4 T Cells. 9, 1–15 (2014).

5. Shi, L. Z. et al. HIF1α–dependent glycolytic pathway orchestrates a metabolic checkpoint for the differentiation of T_H_ 17 and T_reg_ cells. J. Exp. Med. 208, 1367–1376 (2011).

6. Xu, T. et al. Metabolic control of TH17 and induced Treg cell balance by an epigenetic mechanism. Nature (2017). doi:10.1038/nature23475

7. Araki, K. et al. mTOR regulates memory CD8 T-cell differentiation. Nature 460, 108–112 (2009).

8. Pearce, E. L. et al. Enhancing CD8 T-cell memory by modulating fatty acid metabolism. Nature 460, 103–107 (2009).

9. Rao, R. R., Li, Q., Odunsi, K. & Shrikant, P. A. The mTOR kinase determines effector versus memory CD8+ T cell fate by regulating the expression of transcription factors T-bet and Eomesodermin. Immunity 32, 67–78 (2010).

10. Sukumar, M. et al. Inhibiting glycolytic metabolism enhances CD8+ T cell memory and antitumor function.pdf. 123, 4479–4488 (2013).

11. Gubser, P. M. et al. Rapid effector function of memory CD8+T cells requires an immediate-early glycolytic switch. Nat. Immunol. 14, 1064–1072 (2013).

12. Zenhaeusern, G. et al. A high-mobility, low-cost phenotype defines human effector-memory CD8 + T cells. Blood 113, 95–99 (2009).

13. Dimeloe, S. et al. The Immune-Metabolic Basis of Effector Memory CD4+ T Cell Function under Hypoxic Conditions. J. Immunol. 196, 106–14 (2016).

14. Bantug, G. R. et al. Mitochondria-Endoplasmic Reticulum Contact Sites Function as Immunometabolic Hubs that Orchestrate the Rapid Recall Response of Memory CD8 + T Cells. Immunity 48, 542–555.e6 (2018).

15. Chang, C.-H. et al. Posttranscriptional control of T cell effector function by aerobic glycolysis. Cell 153, 1239–51 (2013).

16. Geltink, R. I. K. et al. Article Mitochondrial Priming by CD28. Cell 171, 385–390.e11 (2017).

17. Mestas, J. & Hughes, C. C. W. Of Mice and Not Men: Differences between Mouse and Human Immunology. J. Immunol. 172, 2731–2738 (2004).

18. Beura, L. K. et al. Normalizing the environment recapitulates adult human immune traits in laboratory mice. Nature 532, 512–516 (2016).

19. Akondy, R. S. et al. Origin and differentiation of human memory CD8 T cells after vaccination. Nature 552, 362–367 (2017).

20. Pilyugin, S. S., Ganusov, V. V., Murali-Krishna, K., Ahmed, R. & Antia, R. The rescaling method for quantifying the turnover of cell populations. J. Theor. Biol. 225, 275–283 (2003).

21. Choo, D. K., Murali-Krishna, K., Anita, R. & Ahmed, R. Homeostatic Turnover of Virus-Specific Memory CD8 T Cells Occurs Stochastically and Is Independent of CD4 T Cell Help. J. Immunol. 185, 3436–3444 (2010).

22. Doench, J. G. et al. Optimized sgRNA design to maximize activity and minimize off-target effects of CRISPR-Cas9. Nat. Biotechnol. 34, 184–191 (2016).

23. Alves, T. C. et al. Integrated, Step-Wise, Mass-Isotopomeric Flux Analysis of the TCA Cycle. Cell Metab. 22, 936–947 (2015).

24. van der Windt, G. J. W. & Pearce, E. L. Metabolic switching and fuel choice during T-cell differentiation and memory development. Immunol. Rev. 249, 27–42 (2012).

25. Yao, C.-H. et al. Identifying off-target effects of etomoxir reveals that carnitine palmitoyltransferase I is essential for cancer cell proliferation independent of β-oxidation. PLOS Biol. 16, e2003782 (2018).

26. Jha, A. K. et al. Network integration of parallel metabolic and transcriptional data reveals metabolic modules that regulate macrophage polarization. Immunity 42, 419–430 (2015).

27. Geiger, R. et al. L-Arginine Modulates T Cell Metabolism and Enhances Survival and Anti-tumor Activity. Cell 167, 829–842.e13 (2016).

28. Martin, E. et al. CTP synthase 1 deficiency in humans reveals its central role in lymphocyte proliferation. Nature 510, 288–92 (2014).

29. Angela, M. et al. Fatty acid metabolic reprogramming via mTOR-mediated inductions of PPARγ directs early activation of T cells. Nat. Commun. 7, 1–15 (2016).

30. Salmond, R. J. mTOR Regulation of Glycolytic Metabolism in T Cells. Front. Cell Dev. Biol. 6, 1–9 (2018).

31. Au-Yeung, B. B., Shah, N. H., Shen, L. & Weiss, A. ZAP-70 in Signaling, Biology, and Disease. Annu. Rev. Immunol. 36, 127–156 (2018).

32. Shin, H. J. et al. Rapamycin and IL-2 reduce lethal acute graft-versus-host disease associated with increased expansion of donor type CD4+CD25+Foxp3+regulatory T cells. Blood 118, (2011).

33. Albert, M. H., Yu, X. Z., Martin, P. J. & Anasetti, C. Prevention of lethal acute GVHD with an agonistic CD28 antibody and rapamycin. Blood 105, 1355–1361 (2005).

34. Zhao, H. & Caflisch, A. Discovery of ZAP70 inhibitors by high-throughput docking into a conformation of its kinase domain generated by molecular dynamics. *Bioorganic Med*. Chem. Lett. 23, 5721–5726 (2013).

35. Jacinto, E. et al. Mammalian TOR complex 2 controls the actin cytoskeleton and is rapamycin insensitive. Nat. Cell Biol. 6, 1122–1128 (2004).

36. Oates & Mylari. Aldose reductase inhibitors: therapeutic implications for diabetic complications. Expert Opin. Investig. Drugs 8, 2095–2119 (1999).

37. Johnson, Z. I. et al. TNF-α Promotes Nuclear Enrichment of TonEBP/NFAT5 to Selectively Control Inflammatory but not Osmoregulatory Responses in Nucleus Pulposus Cells. J. Biol. Chem. jbc.M117.790378 (2017). doi:10.1074/jbc.M117.790378

38. Ramana, K. V. & Srivastava, S. K. Mediation of aldose reductase in lipopolysaccharide-induced inflammatory signals in mouse peritoneal macrophages. Cytokine 36, 115–122 (2006).

39. Sánchez-Gómez, F. J. et al. Detoxifying Enzymes at the Cross-Roads of Inflammation, Oxidative Stress, and Drug Hypersensitivity: Role of Glutathione Transferase P1-1 and Aldose Reductase. Front. Pharmacol. 7, 237 (2016).

40. Yadav, U. C. S., Ramana, K. V. & Srivastava, S. K. Aldose reductase inhibition suppresses airway inflammation. Chem. Biol. Interact. 191, 339–345 (2011).

41. Lee, C. F. et al. Preventing Allograft Rejection by Targeting Immune Metabolism. Cell Rep. 13, 760–770 (2015).

42. S., M., V.K., B., K.L.W., S., B.S., S. & D.E., G. Protective effects of glutamine antagonist 6-diazo-5-oxo-L-norleucine in mice with alphavirus encephalomyelitis. J. Virol. 90, 9251–9262 (2016).

43. Angela, M. et al. Fatty acid metabolic reprogramming via mTOR-mediated inductions of PPARγ directs early activation of T cells. Nat. Commun. 7, 13683 (2016).

44. Kanehisa, M. & Goto, S. KEGG: kyoto encyclopedia of genes and genomes. Nucleic Acids Res. 28, 27–30 (2000).

45. Jasmine Chong, Othman Soufan, Carin Li, Iurie Caraus, Shuzhao Li, Guillaume Bourque, David S Wishart, Jianguo Xia; MetaboAnalyst 4.0: towards more transparent and integrative metabolomics analysis, Nucleic Acids Research, Volume 46, Issue W1, 2 July 2018, Pages W486–W494, https://doi.org/10.1093/nar/gky310

46. Alexander Dobin, Carrie A. Davis, Felix Schlesinger, Jorg Drenkow, Chris Zaleski, Sonali Jha, Philippe Batut, Mark Chaisson, Thomas R. Gingeras; STAR: ultrafast universal RNA-seq aligner, Bioinformatics, Volume 29, Issue 1, 1 January 2013, Pages 15–21, https://doi.org/10.1093/bioinformatics/bts635

47. Yang Liao, Gordon K. Smyth, Wei Shi; featureCounts: an efficient general purpose program for assigning sequence reads to genomic features, Bioinformatics, Volume 30, Issue 7, 1 April 2014, Pages 923–930, https://doi.org/10.1093/bioinformatics/btt656

48. Grossman, Robert L., Heath, Allison P., Ferretti, Vincent, Varmus, Harold E., Lowy, Douglas R., Kibbe, Warren A., Staudt, Louis M. (2016) Toward a Shared Vision for Cancer Genomic Data. New England Journal of Medicine 375:12, 1109–1112

49. UCSC Genome Browser: Kent WJ, Sugnet CW, Furey TS, Roskin KM, Pringle TH, Zahler AM, Haussler D. The human genome browser at UCSC. Genome Res. 2002 Jun;12(6):996–1006.

50. Love MI, Huber W, Anders S (2014). “Moderated estimation of fold change and dispersion for RNA-seq data with DESeq2.” Genome Biology, 15, 550. doi: 10.1186/s13059-014-0550-8

51. Huang DW, Sherman BT, Lempicki RA. Bioinformatics enrichment tools: paths toward the comprehensive functional analysis of large gene lists. Nucleic Acids Res. 2009;37(1):1–13.

52. Hänzelmann S, Castelo R, Guinney J (2013). “GSVA: gene set variation analysis for microarray and RNA-Seq data.” BMC Bioinformatics, 14, 7. doi: 10.1186/1471-2105-14-7, http://www.biomedcentral.com/1471-2105/14/7

53. Sergushichev et al. GAM: a web-service for integrated transcriptional and metabolic network analysis. Nucleic Acids Research. http://dx.doi.org/10.1093/nar/gkw266

54. Goenawan, Ivan H., Kenneth Bryan, and David J. Lynn. “DyNet: visualization and analysis of dynamic molecular interaction networks.” Bioinformatics (2016): btw187

55. Haug et al. MetaboLights-- an open-access general-purpose repository for metabolomics studies and associated meta-data. Nucl. Acids Res. (2013) doi: 10.1093/nar/gks1004

