## Supplementary material for "Integrated Metabolomic and Transcriptomic Profiling Reveals Novel Activation-Induced Metabolic Networks in Human T cells": Sup Table 4

| Ratio | Precursor Label | Product Label | Tracer |
|---|---|---|---|
| $\varphi PyrGlu$ | $[U-{}^{13}C_3]Pyruvate$ | $\sum Glutamate_{C4,5^{+2}}$ | [U-$^{13}C_6$]Glucose |
| $\varphi PyrAc$ | $[U-{}^{13}C_3]Pyruvate$ | $[1,2-{}^{13}C_2]Acetyl-CoA$ | [U-$^{13}C_6$]Glucose |
| $\varphi AcCit$ | $[1,2-{}^{13}C_2]Acetyl-CoA$ | $\sum Citrate_{C4,5^{+2}}$ | [U-$^{13}C_6$]Glucose |
| $\varphi CitGlu$ | $\sum Citrate_{C4,5^{+2}}$ | $\sum Glutamate_{C4,5^{+2}}$ | [U-$^{13}C_6$]Glucose |
| $\varphi PyrOAA$ | $[U-{}^{13}C_3]Pyruvate-OAA_{PC}(M+3)$ | $OAA_{PC}(M+3)$ | [U-$^{13}C_6$]Glucose |
| $\varphi GlnGlu$ | $1-(M+0)Glutamine$ | $1-(M+0)Glutamate$ | [1,2-$^{13}C_2$]Glutamine |
| $\varphi GluCit$ | $(M+2)Glutamate$ | $(M+2)Citrate$ | [1,2-$^{13}C_2$]Glutamine |
| $\varphi CitMal$ | $(M+2)Citrate$ | $(M+2)Malate$ | [1,2-$^{13}C_2$]Glutamine |

- $\sum Glutamate_{C4,5^{+2}}$ refers to the sum of all isotopomers of glutamate that have a label in carbons 4 and 5 according to the IUPAC nomenclature.
- $\sum Citrate_{C4,5^{+2}}$ refers to the sum of all isotopomers of citrate that have a label in carbons 4 and 5. The carbon numbering for citrate was defined according to the standard numbering of carbons in glutamate, as detailed by Alves et al.[1]
- $(M+n)MetaboliteName$ refers to a metabolite with n $^{13}C$ carbons and the rest $^{12}C$ carbons
- $OAA_{PC}(M+3)$ refers to portion of M+3 isotopologue of OAA formed directly via Pyruvate Carboxylase (PC), as described by Alves et al.[1]

References:


1. Alves, T. C. et al. Integrated, Step-Wise, Mass-Isotopomeric Flux Analysis of the TCA Cycle. Cell Metab. 22, 936–947 (2015).
